# EWF 2.0: Exact sampling of allele trajectories using the Wright–Fisher diffusion with time-varying demography

**DOI:** 10.1101/2025.10.02.679989

**Authors:** Jaromir Sant, Paul A. Jenkins, Jere Koskela, Dario Spanò

## Abstract

Accurate modelling of allele frequency trajectories requires incorporation of both genetic mechanisms such as selection and mutation, as well as realistic population demography. Accounting for a non-constant demography within a Wright–Fisher diffusion framework induces a time-inhomogenous drift coefficient, a regime falling outside the scope of existing exact simulation routines. To address this gap, we introduce `EWF 2.0`, an exact simulation algorithm that accommodates time-varying demography within Wright–Fisher diffusions whilst retaining all the functionality of previous `EWF` versions. We validate correctness using distributional tests (Kolmogorov–Smirnov, QQ plots), confirming agreement with theoretical expectations. In spite of its greater generality, `EWF 2.0` retains the same runtime as in previous versions, ensuring computational efficiency and scalability. All software is available at https://github.com/JaroSant/EWF. `EWF 2.0` is particularly valuable for bridge simulation, where existing methods cannot handle time-varying mutation and selection rates. For a specified demographic history, mutation parameters, selection function and sampling times, `EWF 2.0` generates exact draws from the law of the corresponding Wright–Fisher diffusion or diffusion bridge.

## 1 Background

Temporal genomic datasets are increasingly used to study how mutation, selection, and genetic drift shape allele frequency trajectories through time (Bollback *et al*., 2008; Malaspinas et al., 2012; Schraiber *et al*., 2016; Ferrer-Admetlla et al., 2016; Sohail et al., 2021). A widely used probabilistic framework for modelling such dynamics is the Wright–Fisher diffusion, which arises as a diffusion limit for a large class of models, including the classical Wright–Fisher model (Ethier and Kurtz (1986, Chapter 10); Karlin and Taylor (1981, Chapter 15); Ewens (2004); Crow and Kimura (1970)), and therefore provides a robust description of allele-frequency dynamics that is not tied to any single microscopic reproduction mechanism. The Wright–Fisher diffusion is particularly valued for its flexibility in modelling the combined effects of mutation, selection, and genetic drift on allele frequency trajectories, with many inference methods in population genetics building on this process to analyse temporal variation in genetic data (Bollback *et al*., 2008; Malaspinas et al., 2012; Schraiber et al., 2016; Ferrer-Admetlla et al., 2016; Sohail et al., 2021; He *et al*., 2020a).

Despite its widespread use, working with the Wright–Fisher diffusion remains difficult in practice. Although an explicit infinite-series representation of the transition density has long been available (Griffiths, 1979; Tavaré, 1984), it is computationally unwieldy, forcing most practical methods to rely on approximations or discretisation schemes, thereby introducing biases that are hard to quantify. These difficulties become even more pronounced when population size varies through time, since changes in effective population size alter the strength of mutation, selection, and genetic drift and therefore substantially affect allele frequency trajectories. If such demographic variation is ignored, its effects may be confounded with changes in mutation or selection pressures. Frameworks that accommodate non-constant demographies are now common, with spectral methods providing pointwise evaluations of the transition density under varying effective population sizes (Steinrücken *et al*., 2015), and inferential approaches directly incorporating demographic change into the methodology (Schraiber *et al*., 2016; He *et al*., 2020b). Related approaches have also been considered in the context of exact conditioned (bridge) simulation for finite Wright–Fisher models (Zhao *et al*., 2014). Despite being exact at the level of the discrete model, such a scheme requires explicit simulation of every generation. In the biologically relevant regime of large populations evolving over long timescales, this may involve hundreds of thousands of generations. By contrast, the diffusion formulation permits direct simulation on evolutionary timescales, leading to substantial computational savings in neutral and near-neutral settings (see Appendix 8). However, to date no software exists to permit *exact* simulation under a time-varying demography.

Here and throughout, we take “exact” to mean that trajectories are sampled directly from the target Wright–Fisher diffusion law itself, rather than from a time-discretised or other approximation of the model. Thus in an implementation of an exact simulation method the only sources of error are those arising from floating point arithmetic and from model error, i.e. the discrepancy between a model and the physical phenomena it seeks to describe (an issue inherent to any mathematical model). For the rest of the paper we do not address model error any further; the Wright–Fisher diffusion is a very well-established model in population genetics and its ability to capture the salient features of real allele frequency trajectories has been studied extensively (Crow and Kimura, 1970; Kimura, 1983; Ewens, 2004).

Exact simulation is particularly useful as a backbone for simulation-based inference frameworks and related inferential tools Ferrer-Admetlla et al. (2016); Foll et al. (2015); Schraiber et al. (2016); He et al. (2020a), as well as for benchmarking approximate methods Bollback et al. (2008); Sohail et al. (2021, 2022); Xu et al. (2023), generating realistic simulated datasets, and studying scenarios where discretisation error may become non-negligible, such as strong selection or rapidly changing demography. The aim of this article is to provide an exact simulation tool for use within downstream genetic analyses in the presence of non-constant demographies. We emphasise, however, that the present work is concerned exclusively with simulation and does not perform inference.

A major step forward in exact simulation was the introduction of an exact sampling technique for Wright–Fisher diffusions by Jenkins and Spanò (2017), which was subsequently implemented as `EWF` in Sant et al. (2023). This enables efficient simulation across a wide class of Wright–Fisher diffusion models, including non-neutral diffusions and conditioned trajectories, but only under the assumption of constant population size. That assumption is restrictive, since demographic variation is ubiquitous in natural populations and strongly influences the behaviour of allele frequency trajectories.

The main aim of this work is to extend this class of exact simulation schemes to accommodate piece-wise constant demographies, presenting both the mathematical framework and its implementation in `EWF 2.0`. Piecewise constant demographic histories naturally capture common scenarios such as bottlenecks and population expansions, whilst also providing approximations to more general continuously varying demographies. In practice, piecewise constant demographic histories are commonly inferred from genome-wide data using coalescent-based methods Li and Durbin (2011); Palamara et al. (2018); Nait Saada et al. (2020), with changepoints corresponding to the inferred boundaries of demographic epochs. Finer discretisations provide closer approximations to continuously varying demographic histories, at the cost of introducing a larger number of epochs. Each change in effective population size induces a corresponding change in the diffusion-scaled mutation and selection parameters, leading to a time-inhomogeneous Wright–Fisher diffusion and consequently requiring non-trivial modifications to existing exact simulation schemes.

Despite these modifications, `EWF 2.0` maintains the asymptotic runtime behaviour of the original `EWF` algorithm Sant et al. (2023). In practice, `EWF 2.0` exhibits marginally faster runtimes compared to the original implementation, since the additional overhead costs of time-varying parameters are offset by gains arising from implementation-level caching and storage optimisations.

The software can simulate allele frequency trajectories conditioned on their endpoint, known as a diffusion *bridge*. Bridge diffusions arise naturally when allele frequencies are observed at a finite number of time points and one wishes to do statistical inference conditional those observations. This setting is common in ancient DNA studies, experimental evolution, and other temporal population-genetic datasets, where genetic samples are available only at discrete and often sparse time points Bollback et al. (2008); Malaspinas et al. (2012); Schraiber et al. (2016); Ferrer-Admetlla et al. (2016); Sohail et al. (2021, 2022); He et al. (2020b,a). Beyond providing biologically realistic trajectory reconstructions, bridge simulation also plays an important role in genomic selection analyses: when loci are pre-screened on the basis of large observed allele-frequency shifts, ascertainment bias may be introduced. Conditioning on the observed endpoints through bridge simulation provides a principled way of correcting for this effect, thereby avoiding spurious inflation of mutation or selection estimates (Nielsen and Signorovitch, 2003; Foll *et al*., 2015).

`EWF 2.0` enables exact sampling of Wright–Fisher diffusions and bridges when mutation and selection parameters differ across demographic epochs, while preserving all features of the original `EWF` package, including boundary absorption/reflection and arbitrary frequency-dependent selection. This substantially broadens the scope of exact Wright–Fisher simulation and strengthens its utility for modern population-genetic inference. A comparison of `EWF 2.0` with some related software packages is shown in Table 1.

**Table 1:** Comparison of existing Wright–Fisher methods for both simulation and likelihood evaluation: “WF model” is the method found in Zhao et al. (2013), “WF bridge” is the method in Zhao et al. (2014), `SpectralTDF` is the method found in Steinrücken et al. (2015), and `EWF` is the one in Sant et al. (2023). Here “Exact” refers to exact simulation from the target diffusion law rather than from a time-discretized approximation.

| Method | Exact | Variable<br>$N_e(t)$ | Bridge<br>simulation | Selection | Trajectory<br>simulation | Wright–Fisher<br>diffusion |
| --- | --- | --- | --- | --- | --- | --- |
| WF model | × | ✓ | × | ✓ | × | × |
| WF bridge | ✓ | × | ✓ | ✓ | ✓ | × |
| **SpectralTDF** | × | ✓ | × | ✓ | × | ✓ |
| **EWF** | ✓ | × | ✓ | ✓ | ✓ | ✓ |
| **EWF 2.0** | ✓ | ✓ | ✓ | ✓ | ✓ | ✓ |

The rest of this article is organised as follows: Section 2 details the mathematical setup and implementation of the method, Section 3 illustrates its performance, and Section 4 concludes with a brief discussion. Appendix 1 derives the various possible bridge diffusion transition densities, with Appendix 2 providing the blueprint for how exact simulation can be achieved in one particular case. Appendix 3 then extends these arguments to the remaining cases, with details regarding the small time approximations found in Appendix 4. Full details on the output verification and tests can be found in Appendix 5, whilst details on non-neutral simulations can be found in Appendix 6. Appendix 7 provides further details on the horse demography used in Figure 2, and Appendix 8 details the computational advantages of the diffusion formulation over a discrete model.

## 2 Implementation

We model the frequency of an allele evolving under mutation, selection, and genetic drift in a population with time-varying effective population size through the Wright–Fisher diffusion (Ethier and Kurtz, 1986; Karlin and Taylor, 1981; Ewens, 2004). In this setting, the stochastic process *X* := (*X*_*t*_)_*t*≥0_ describes the allele frequency trajectory through time, with *X*_*t*_ ∈ [0, 1] representing the population frequency of the allele at time *t*, satisfying the stochastic differential equation (Ethier and Kurtz (1986, Chapter 10); Karlin and Taylor (1981, Chapter 15); Ewens (2004))

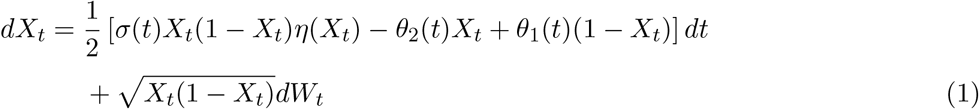

for *t* ≥ 0 with *X*_0_ ∈ [0, 1]. Here the drift term models the effects of mutation and selection through the time-varying mutation parameter ***θ***(*t*) = (*θ*_1_(*t*), *θ*_2_(*t*)), the selection coefficient *σ*(*t*), and the frequency-dependent selection function *η*(*x*), whilst the diffusion term captures genetic drift. We further assume that the selection function is of the form 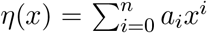 for some finite *n*, which encompasses a wide class of selective regimes including genic selection (*η*(*x*) ≡ 1) and diploid selection (*η*(*x*) = *h* + *x*(1 − 2*h*) with *h* the dominance parameter).

The effective population size is assumed to be given by a known piecewise constant function *N*_*e*_(*t*). Changes in *N*_*e*_(*t*) alter the strength of genetic drift and induce corresponding changes in the diffusion-scaled mutation and selection parameters via *θ*_*j*_(*t*) = 2*N*_*e*_(*t*)*µ*_*j*_ for *j* = 1, 2, and *σ*(*t*) = 2*N*_*e*_(*t*)*γ*, where *µ*_*j*_ denotes the corresponding mutation rate per generation and *γ* denotes the selection coefficient. We refer to each interval over which *N*_*e*_(*t*) is constant as an *epoch*. Equation (1) is written in diffusion time units, obtained after the usual Wright–Fisher diffusion rescaling applied within each epoch; the corresponding parameters are therefore diffusion-scaled quantities. Throughout, time is measured in diffusion units unless otherwise stated. Within a demographic epoch having effective population size *N*_*e*_(*t*), the corresponding real time *τ* measured in years satisfies

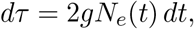

where *g* denotes the generation time in years, and *t* denotes diffusion time. Thus conversion between diffusion time and real time is obtained by integrating the effective population size over the corresponding demographic epoch. In particular, when *N*_*e*_(*t*) is constant over an epoch, diffusion time scales linearly with the number of generations elapsed.

For illustration, consider two epochs (−∞, *s*) and [*s*, ∞) for some *s >* 0, with corresponding parameters ***θ***, *σ* and ***θ***^***′***^, *σ*^′^ over the two intervals, corresponding to two different effective population sizes *N*_*e*_ and 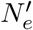. Suppose the diffusion starts at *x* ∈ (0, 1) at time 0 and ends at *z* ∈ (0, 1) at time *t > s*. In the neutral case (i.e. *σ* = 0 = *σ*^′^) with strictly positive mutation parameters, the bridge transition density at changepoint time *s* admits the mixture representation (Jenkins and Spanò, 2017, Proposition 2)

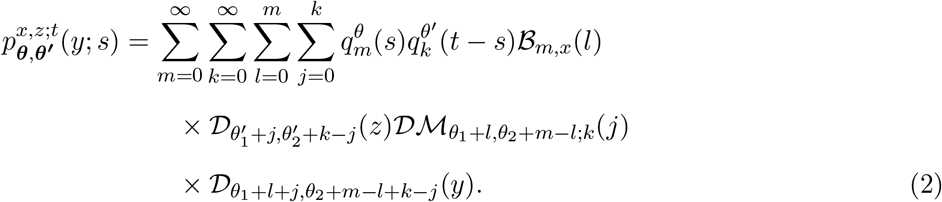

Here 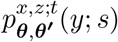 denotes the conditional density of the allele frequency at time *s*, given that the trajectory starts at frequency *x* at time 0 and is conditioned to end at frequency *z* at time *t*. For any *u >* 0, the quantities 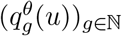 define a probability distribution on the integers, whilst ℬ, D and Dℳ denote the binomial, beta, and Dirichlet-multinomial probability mass functions respectively. Full expressions and notation are provided in Appendix 1.

Equation (2) expresses the bridge density as an infinite mixture of Beta distributions, and is central to the simulation scheme. By extending the alternating-series method of Jenkins and Spanò (2017) to scenarios where mutation parameters differ across demographic epochs, exact sampling from the bridge distribution without time or state space discretisation is possible. In particular, exact draws from (2) can then be generated through the following procedure:

1. Draw (*M, K, L, J* ) from the corresponding mixture distribution

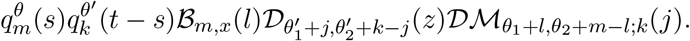
2. Conditional on (*M, K, L, J* ) = (*m, k, l, j*), draw

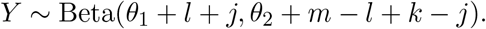

The second step is straightforward, whereas the first requires sampling from an infinite mixture distribution. The precise mathematical statement together with the corresponding proofs are provided in Appendix 2 as well as Appendix 3.

Expressions for the bridge transition density for all combinations of starting and ending frequencies, including the boundary cases *x, z* ∈ {0, 1}, are given in Appendix 1. These cases are especially relevant for *de novo* mutations, where alleles arise at effectively zero frequency before spreading through the population. Since the beta densities in Equation (2) may become singular at the boundaries, these cases require a separate treatment rather than a naive evaluation of the mixture representation.

Our method further allows for sampling bridge trajectories at arbitrary times. Suppose we are interested in generating allele frequencies at time *τ* from the law of a diffusion bridge going from *x* to *z* over a time interval *τ* + *s* + *t* where *τ, s, t >* 0, and *τ* + *s* denotes a time at which we have a demographic changepoint such that the mutation parameter changes from ***θ*** to ***θ***^***′***^ at time *τ* + *s*. Then the joint density of bridge points *y, w* drawn at times *τ* and *τ* + *s* is given by

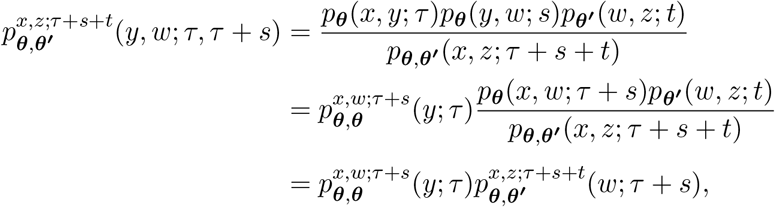

where we have implicitly accounted for changes in *N*_*e*_(*t*) via the mutation parameter. Thus diffusion bridge samples at time *τ* can be generated by integrating out the corresponding samples generated at time *τ* + *s*.

### 2.1 Non-neutral simulation

The algorithm of Jenkins and Spanò (2017, Section 5) for simulating non-neutral Wright–Fisher bridges extends naturally to the setting with changing effective population size. The key idea is to employ a rejection sampler with neutral trajectories as proposals, allowing exact samples from the neutral process to be thinned into exact samples under selection; see Appendix 6 for further details. We note that this sampling approach requires, on average, a number of proposals that grows exponentially with both the time increment and the strength of selection (Jenkins and Spanò, 2017, Proposition 7), rendering it increasingly inefficient for large values of either quantity. Further details on the computational cost of such a scheme can be found in (Jenkins and Spanò, 2017, Section 5).

### 2.2 Small time approximations

For very short intervals *s* or *t* − *s*, the quantities 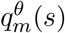 and 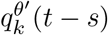 appearing in Equation (2) become numerically unstable (Griffiths, 1984; Jenkins and Spanò, 2017). In this regime we instead approximate the transition by a Gaussian distribution using a central limit theorem of Griffiths (1984, Theorem 4). This approximation is used whenever the interval length falls below a user-defined threshold which is set to 0.1 by default, and which is independent of both demography and the value of the starting point *x* (see (Sant *et al*., 2023, Section 6, Supplementary Material) for details on this choice). Further details on the implementation are provided in Appendix 4, whilst details pertaining to the approximation accuracy can be found in top right panel of Figure 4 as well as Griffiths (1984) and Sant et al. (2023).

## 3 Results

We implemented our method in C++ with a Python front end, building upon the software `EWF`. The relevant sampling routines were suitably modified to allow for non-constant demographies whilst retaining all previous functionality. The package includes example scripts for integrating `EWF 2.0` into population-genetic workflows.

To gauge the implementation’s accuracy and correctness, we generated 10,000 samples of the trajectory of a diffusion for a range of sampling time and parameter regimes, and numerically computed the corresponding transition density by suitably truncating the infinite sums in (2). Across all tested setups, both QQ plots and Kolmogorov–Smirnov tests showed close agreement between simulated and theoretical distributions (see Appendix 5), confirming the correctness of our implementation. The diffusion formulation also offers substantial computational savings relative to direct finite-generation simulation in the neutral case, as illustrated in Appendix 8.

Figure 1 contrasts bridge realisations under a constant and a repeated bottleneck demography, showing how relative to constant demographies the varying demographic model induces noisier allele frequency trajectories during bottleneck periods in line with theory. Accounting for and explicitly modelling these shifts is essential to avoid confounding demographic effects with other genetic forces, underscoring the practical value of `EWF 2.0`.

**Figure 1:**
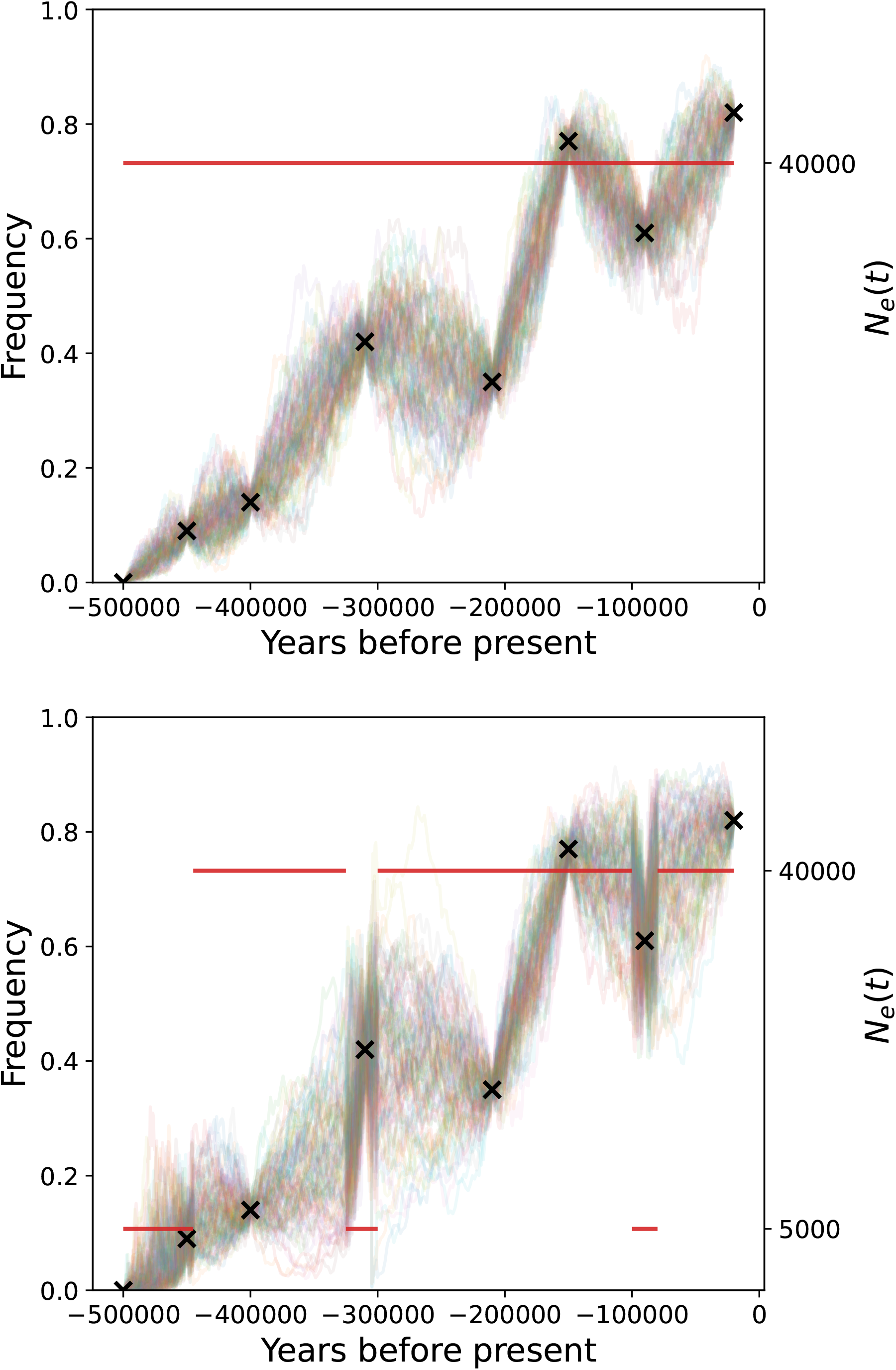
Bridge diffusion samples generated under a constant *N*_*e*_ = 40000 demography (top) and under a repeated bottleneck demography (bottom). True observations are denoted by black crosses, with the corresponding demographies plotted in red. Note that all trajectories pass directly through the black crosses as we are assuming exact observations for illustrative purposes.

**Figure 2:**
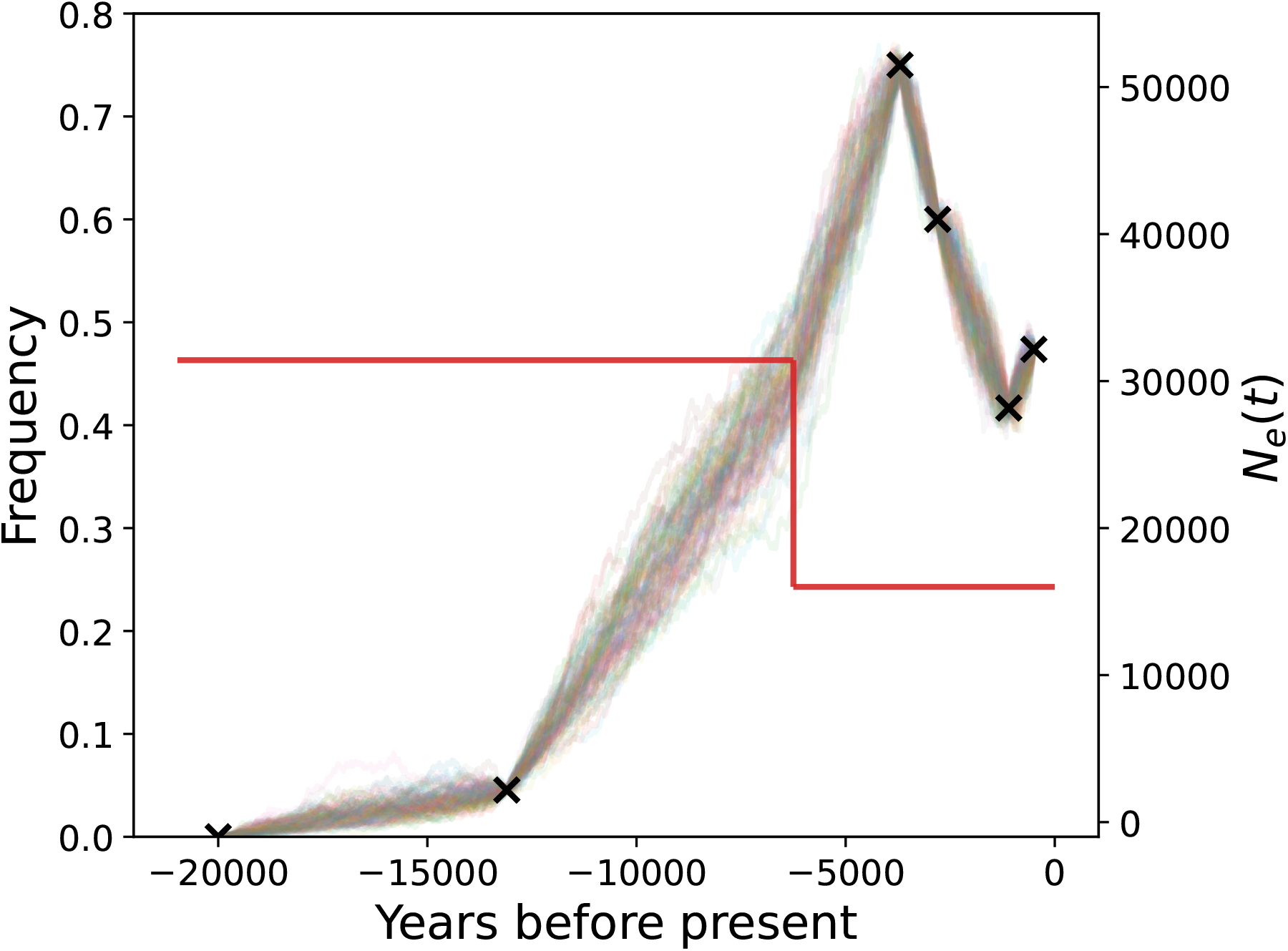
100 candidate trajectories for the horse coat colouration data found in Ludwig et al. (2009) simulated using `EWF 2.0`, using the demography inferred in Der Sarkissian et al. (2015). The observed frequencies (black crosses) as well as the coloured trajectories are assumed to be exact observations of the underlying diffusion for illustrative purposes. Note how the above simulated paths differ from those in Figure 1 in Sant et al. (2023), in particular how the kink in the paths observed here at 6250 years before present is absent there.

To further illustrate the practical use of `EWF 2.0` in simulating data under realistic demographies and plausible parameter regimes, we simulated 100 candidate allele-frequency trajectories for the horse coat coloration dataset (Ludwig *et al*., 2009), using demography inferred in (Der Sarkissian *et al*., 2015, Figure 1B) (see Appendix 7 for full details). We further assumed a generation time of five years as well as biologically plausible per generation mutation and selection parameters of *µ*_1_ = *µ*_2_ = 6.2 × 10^−9^ and *γ* = 7 × 10^−4^ respectively (Lee *et al*., 2014; Ludwig *et al*., 2009). For simplicity, we treat the observed frequencies as direct observations of the underlying latent diffusion, illustrating the utility of our method to simulate trajectories from non-neutral Wright–Fisher diffusions with time-changing demography. Incorporating additional sampling noise, for example through a binomial observation model, can be handled straightforwardly as a separate post-processing step.

## 4 Conclusions

Demographic changes alter the balance of evolutionary forces shaping populations, and accurate models of allele frequency evolution must explicitly account for these dynamics. Here we have shown how exact simulation of allele frequency trajectories using the Wright–Fisher diffusion can be extended to settings with variable effective population size. Our software `EWF 2.0` generalizes the framework of Sant *et al*. (2023) by enabling exact draws from diffusion and diffusion bridge trajectories even when mutation and selection parameters differ across epochs. This is especially important when considering diffusion bridges, where ascertainment bias is appropriately corrected for by conditioning on observed endpoints under realistic demographic histories.

Beyond its theoretical contribution, `EWF 2.0` provides a practical tool for population genetic simulation. It can be readily integrated into pipelines for inference from time-series data, benchmarking of approximate algorithms, and exploration of evolutionary scenarios under demographic change. With a Python interface and efficient C++ routines, it is accessible for use in both methodological and empirical studies, and is particularly timely given the increasing availability of temporal genomic datasets across diverse species (Ludwig *et al*., 2009; Wutke et al., 2016; Loog et al., 2017; Fages et al., 2019; Akbari et al., 2024; Mallick et al., 2024). Its chief limitations are that the proposed method employs an approximation for small time increments owing to numerical instabilities, whilst the non-neutral method suffers from exponentially growing run-times due to the rejection sampling algorithm used. We aim to address these in future work.

## Supplementary Information

## 1 Expressions for the bridge transition density

Consider a Wright–Fisher diffusion bridge started from *x* ∈ (0, 1) at time 0 and ending at *z* ∈ (0, 1) at time *t*, having mutation parameter ***θ*** over the time interval [0, *s*) and ***θ***^***′***^ over [*s, s* + *t*) for some *s* ∈ (0, *t*). The density (with respect to Lebesgue measure) of a point *y* sampled at time *s* is given (Jenkins and Spanò, 2017, Proposition 2) by

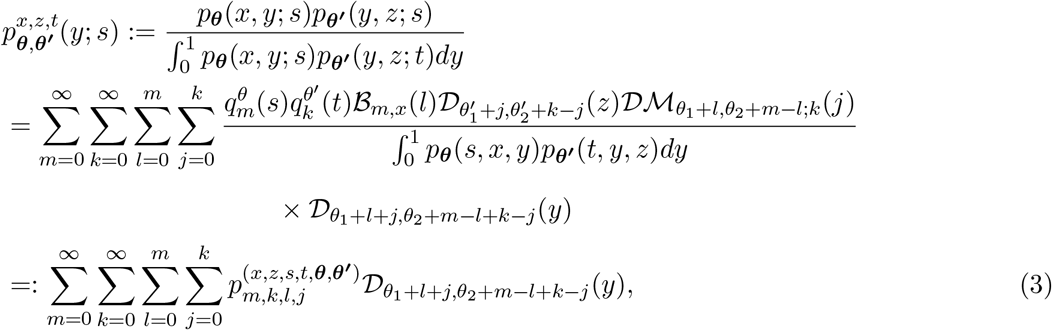

where *θ* := *θ*_1_ + *θ*_2_, *p*_***θ***_(·, ·; ·) denotes the transition density for a Wright–Fisher diffusion with mutation parameter ***θ***,

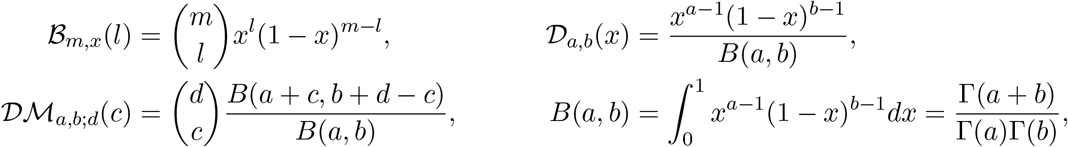

denote the usual binomial probability mass function, beta density, beta-binomial probability mass function and beta function respectively, and Γ(*x*) denotes the gamma function. We observe further that

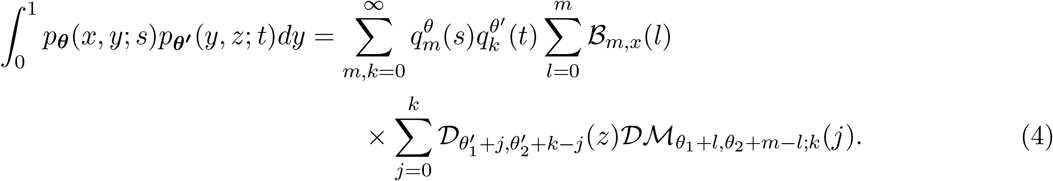

The cases corresponding to when either of *x* or *z* are at the boundary {0, 1} can be derived from (3) by taking the corresponding limit as *x* or *z* go to 0 or 1 in both the denominator and numerator. We illustrate the calculations for the case *x* = 0 and *z* = 1; all other cases follow using similar suitably modified arguments. Observe that if *x* = 0, the only non-zero term in the sum over *l* in (3) and (4) corresponds to the case when *l* = 0, such that

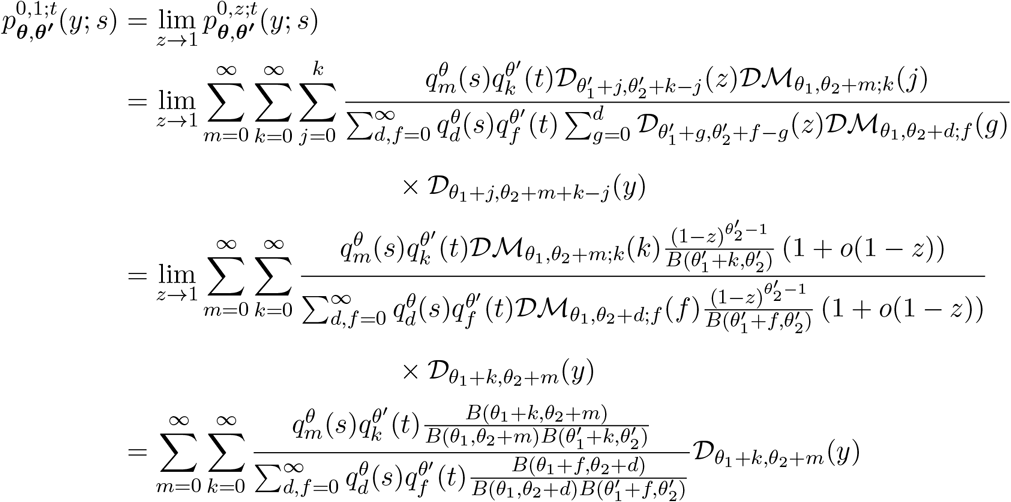

All possible bridge transition densities can be found below. Note that in 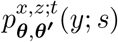, the superscripts *x, z* denote the bridge endpoints; writing *x, z* means that *x, z* ∈ (0, 1), whereas boundary endpoints are written explicitly as 0 or 1. In the next section we show that we can obtain monotonically converging sequences of upper and lower bounds, as in Jenkins and Spanò (2017) for the case when *x, z* ∈ (0, 1), whereas the edge cases when either of *x, z* ∈ {0, 1} are tackled in Section 3, and follow by adapting similar arguments.

We point out that here we are only interested in cases when both ***θ*** and ***θ***^***′***^ have strictly positive entries, since in practice changes in the diffusion-rescaled mutation parameters are driven by corresponding changes in the effective population size rather than in the pre-limiting mutation rate itself. Thus whilst it is theoretically possible to have ***θ*** = (*θ*_1_, *θ*_2_) with *θ*_1_, *θ*_2_ *>* 0 and ***θ***^***′***^ = **0**, we shall not consider such scenarios as this would correspond to the case where *N*_*e*_(*t*) = 0 for some time *t* (i.e. a population collapse, in which case a diffusion approximation is ill-suited). Nonetheless similar calculations to those presented here and in Sant et al. (2023) still hold, and thus exact simulation is still possible.

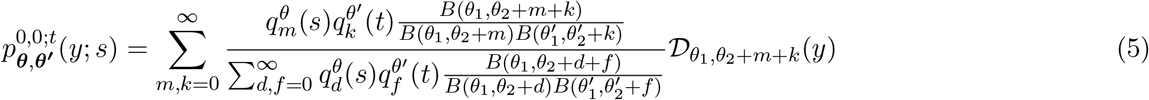

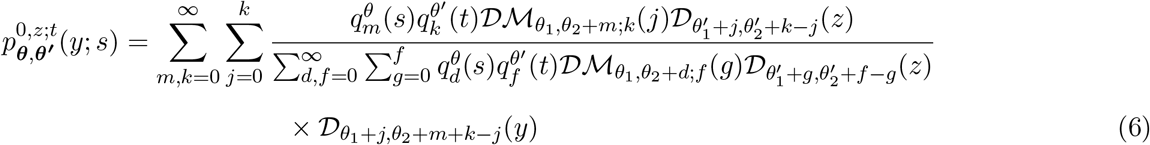

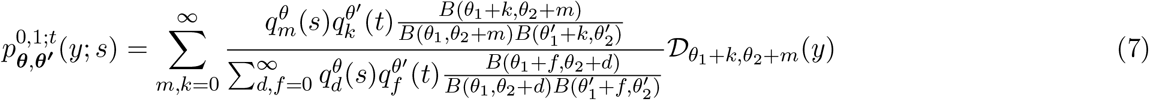

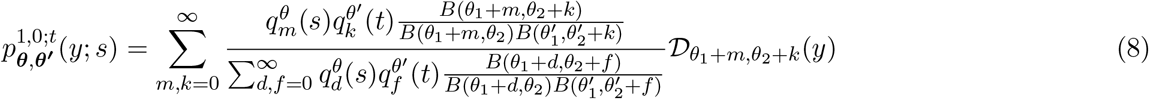

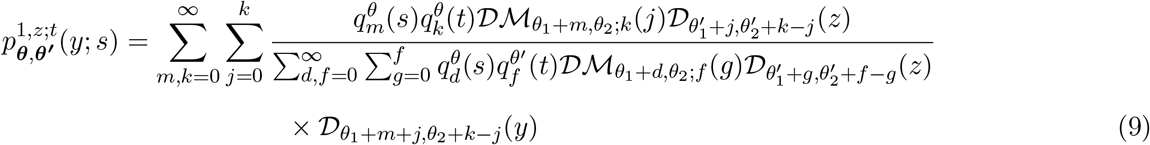

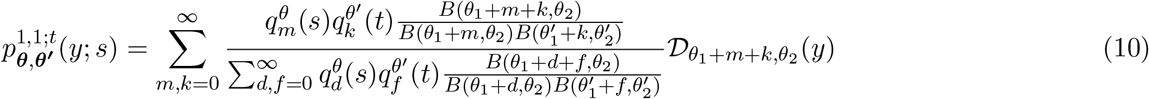

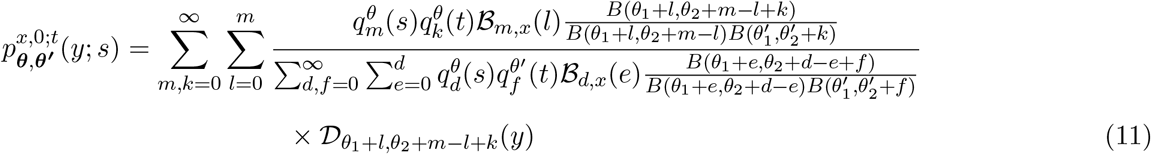

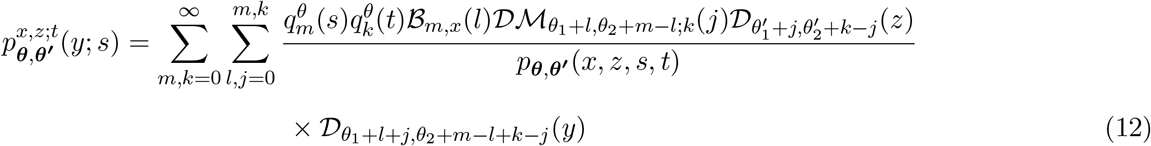

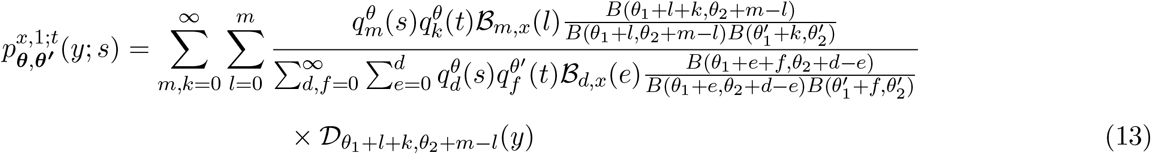

## 2 Simulating exact draws from the density (12)

We can adopt the same approach as in Proposition 4 in Jenkins and Spanò (2017) to obtain monotonically converging sequences of upper and lower bounds for the numerator in (12), however the denominator differs significantly from the one present in Jenkins and Spanò (2017). Here we show how such bounds can be obtained in the case when *x, z* ∈ (0, 1) via Proposition 1 below, whilst in Section 3 we extend these arguments to the remaining cases.

As in (Jenkins and Spanò, 2017, Section 3), these monotonically converging sequences of upper and lower bounds then allow us to generate draws (*m, k, l, j*) ∈ N^4^ (or a suitable subspace of N^4^), from which we can obtain samples from the corresponding bridge diffusion by considering an appropriate draw from a beta distribution.

In view of the infinite series expansion (Griffiths, 1979; Tavaré, 1984)

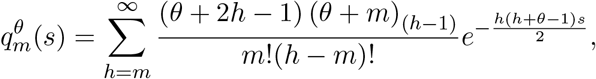

observe that we can re-write

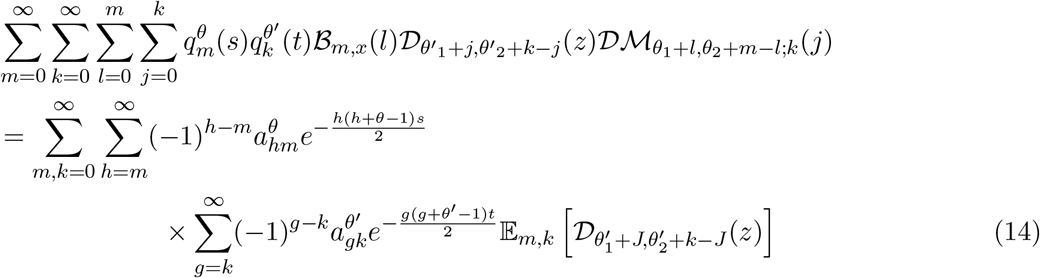

where 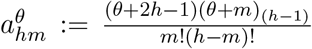, with *a* _(*n*)_: = Γ(*a* + *n*)*/*Γ(*a*) for *a >* 0 and *n* ≥ −1, and the expectation is with respect to the pair of random variables (*L, J* ) such that *L* ∼ Bin(*m, x*) and *J* |*L* = *l* ∼ BetaBin(*k, θ*_1_ + *l, θ*_2_ + *m* − *l*).

Define *λ* := *m* + *k* and *γ* := *h* + *g*, then combining (4) with (14) allows us to deduce that

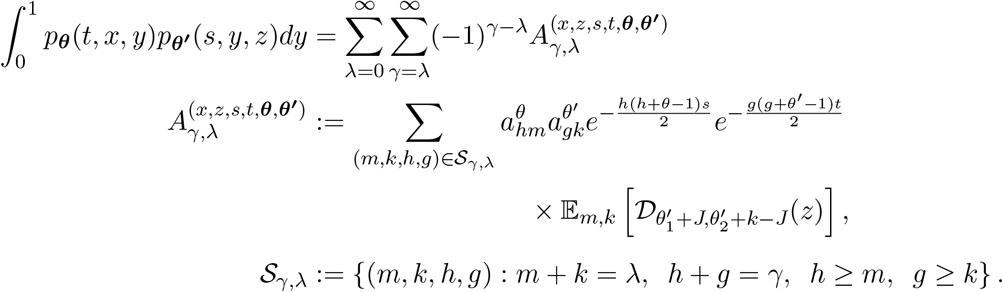

In what follows, we shall drop the superscript in 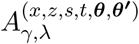 for ease of exposition.

We further define 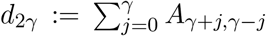 and 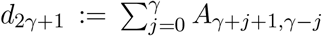, and observe that (4) can be re-written as the alternating series,

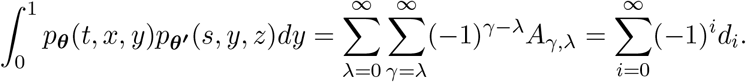

Thus we can in principle adapt Proposition 3 in Jenkins and Spanò (2017) to target this quantity through monotonically converging sequences of upper and lower bounds, namely we want to show that

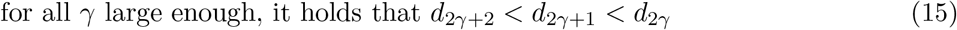

Because of a potential change in mutation parameters at time *t*, adapting this Proposition is not straight forward since we lose the ability to ‘group’ terms in a convenient way.

Instead, for a given *ε* ∈ (0, 1) fixed, define

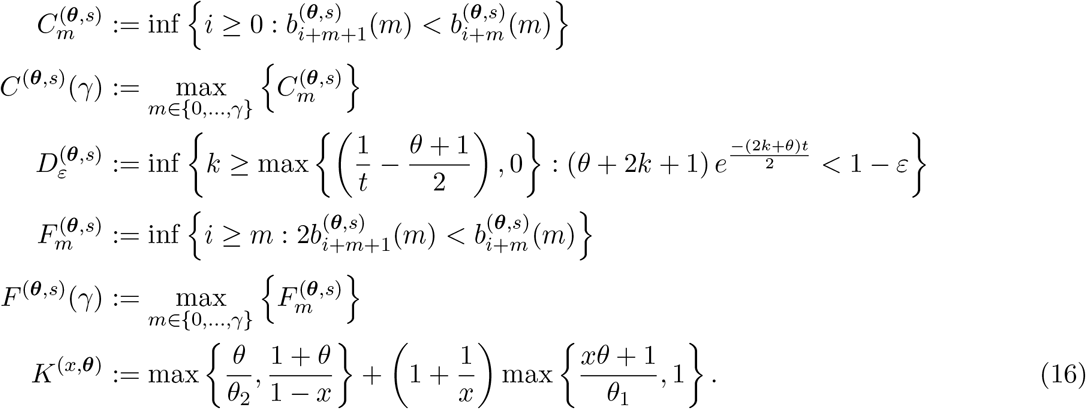

Then we have the following result:

### Proposition 1.

*Let A*_*γ,λ*_, *d*_2*γ*_, *d*_2*γ*+1_,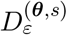, *F* ^(***θ***,*s*)^(*γ*), *K*^(*x*,***θ***)^ *be defined as above, and let ε* ∈ (0, 1) *be some constant. For*

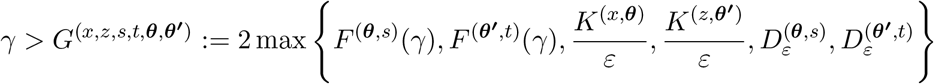

*we have that d*_2*γ*+2_ *< d*_2*γ*+1_ *< d*_2*γ*_.

*Proof*. We start by showing that *d*_2*γ*+1_ *< d*_2*γ*_. For this to be the case, we can expand out the *A*_*λ,γ*_ terms to deduce that

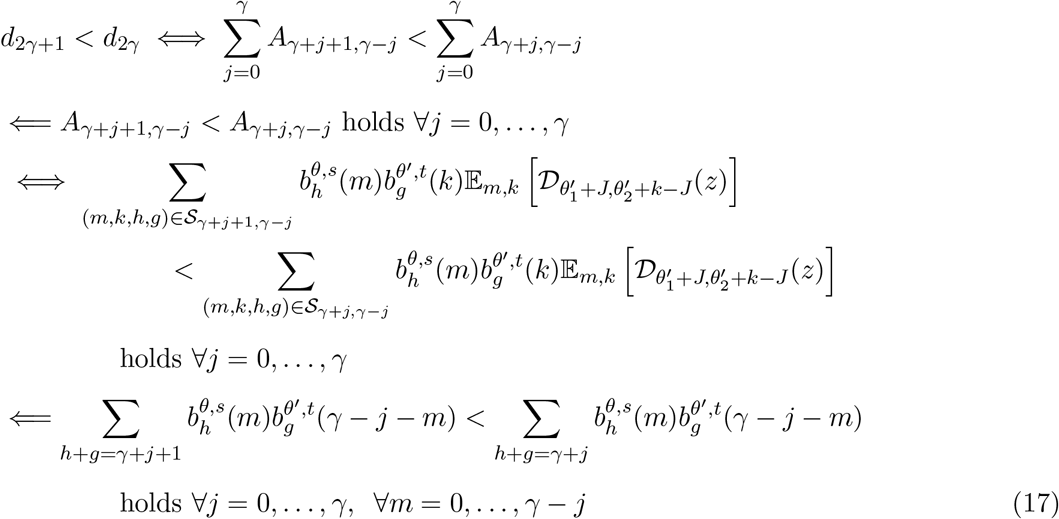

where 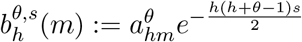.

Consider for the moment a fixed *m* and *j*, and assume that *γ* + *j* + 1 is even (if not similar calculations hold, with suitably altered indexing). We can re-write

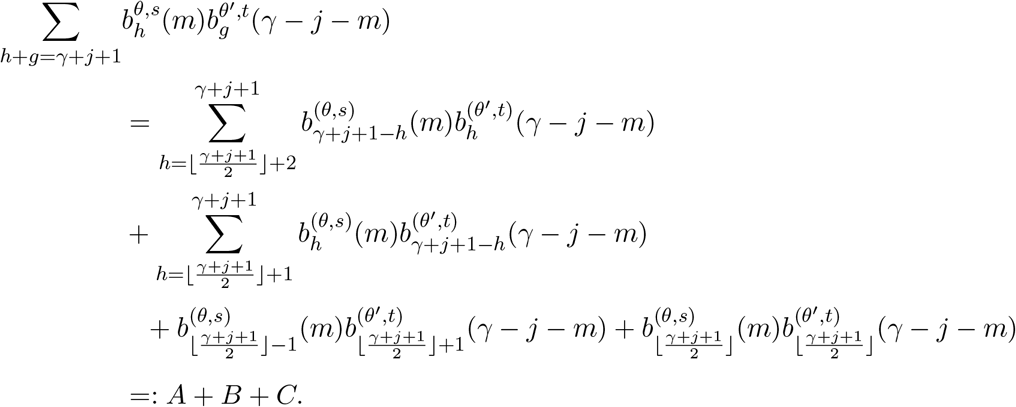

We deal with each of these three terms in turn.

*Term C:*

Recall that for any *k, m* we have that

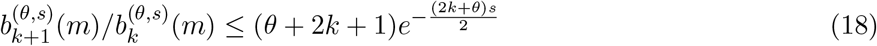

(see (26) and (27) in Jenkins and Spanò (2017)), from which we get that 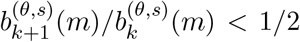 for 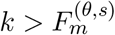. In particular, if *γ* is such that 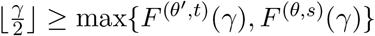, then

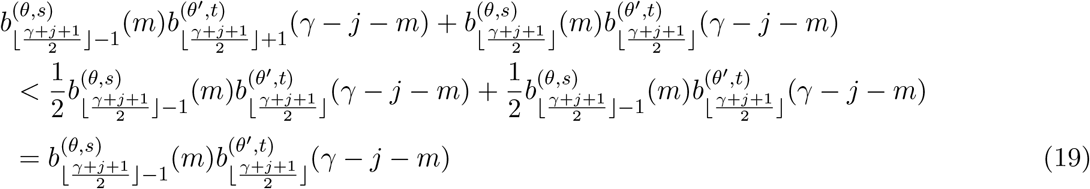

holds independently of *m* and *j*. Recall that if 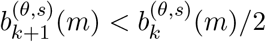 holds for some *k*, it must also hold for all values greater than *k* (see display before eq (27) in Jenkins and Spanò (2017)).

*Term B:*

If *γ* is such that 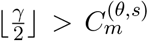, then 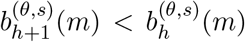 for 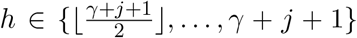. Recall we have fixed *m* and *j* here, but we require (17) to hold ∀*m* = 0, …, *γ* − *j* and ∀*j* = 0, …, *γ*. Thus if we choose 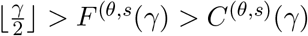, we get that

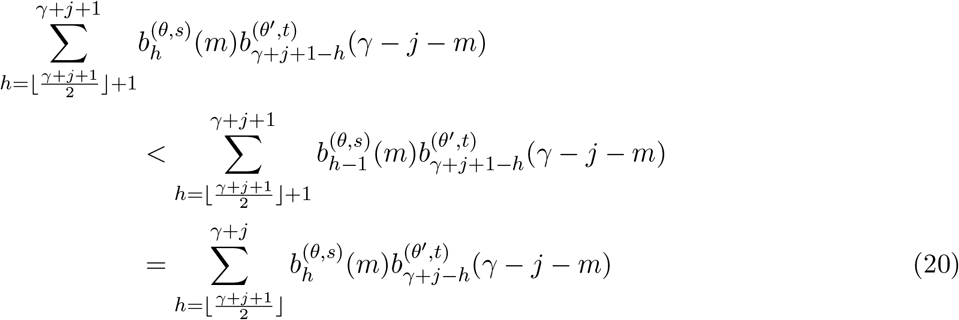

*Term A:*

We can apply a similar argument to the above, namely choose *γ* such that 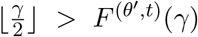, then 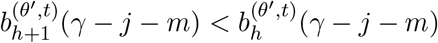 holds for all 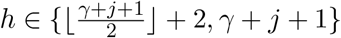, and thus

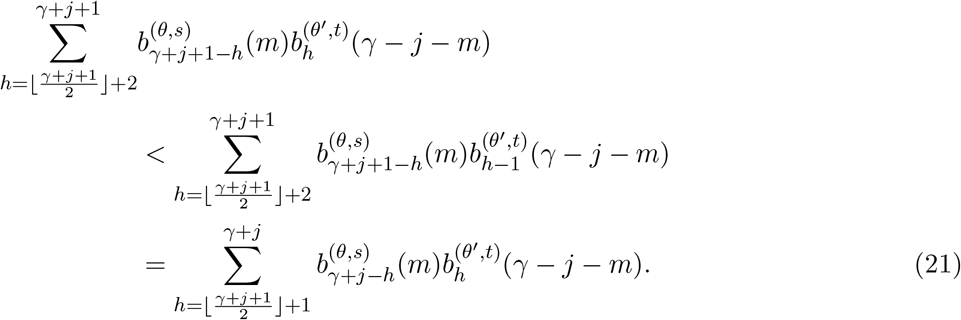

We point out that since 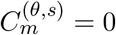 for all 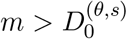 as per Proposition 1(iii) in Jenkins and Spanò (2017), the worst case scenario would be taking 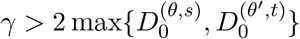 to ensure both (20) and (21) hold.

So if we choose *γ* such that *γ >* 2 max{*F* ^(*θ,s*)^(*γ*), *F* ^(*θ′,t*)^ (*γ*)}, we get that we can combine (19), (20), and (21) to deduce that

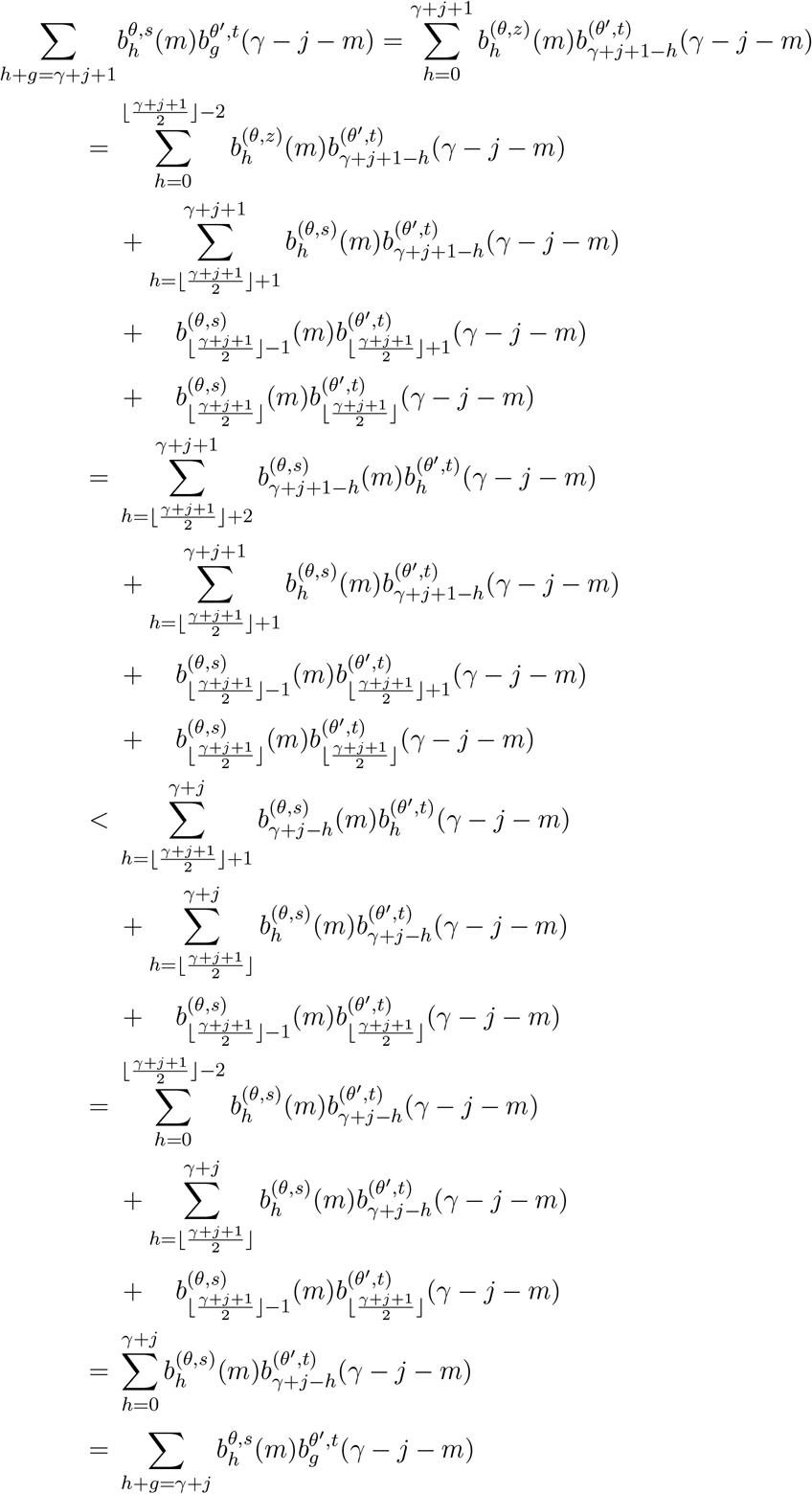

and thus *d*_2*γ*+1_ *< d*_2*γ*_ holds from (17).

It now remains to show that for sufficiently large *γ* we have that *d*_2*γ*+2_ *< d*_2*γ*+1_. Again we use a similar approach to that in the proof of Proposition 3 in Jenkins and Spanò (2017), namely because we have shown that

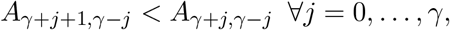

we can increment the first index by 1 and sum over *j* = 1, …, *γ* to deduce

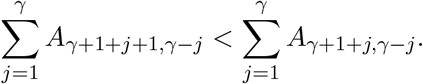

Thus it remains to show that

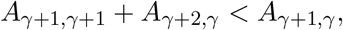

for then it follows that *d* _2*γ*+2_ *< d*_2*γ*+1_ . Observe that if 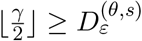, then

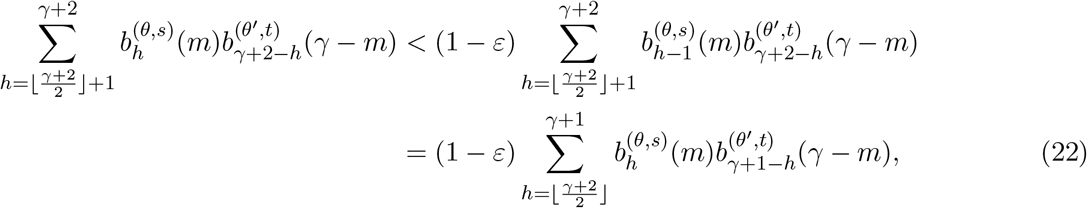

following the same reasoning as in the display above equation (34) in Jenkins and Spanò (2017). We can repeat this argument with 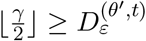 to get

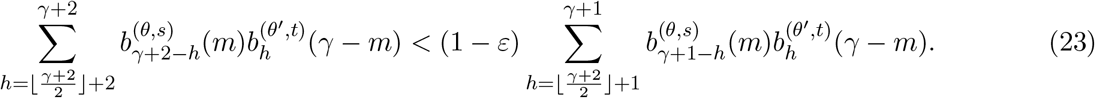

Finally, for 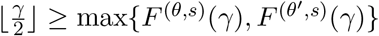, using the same reasoning that led to (19), it follows that

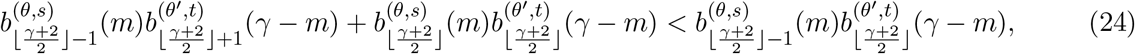

and thus summing (22), (23) and (24) gives

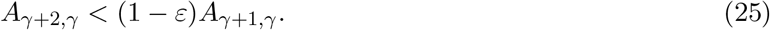

In view of (25), it suffices to show that

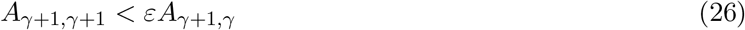

To this end, observe that

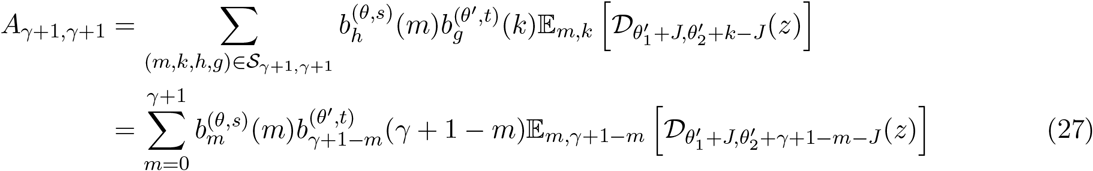

since *γ* + 1 − *h* = *g* ≥ *k* = *γ* + 1 − *m*, and similarly

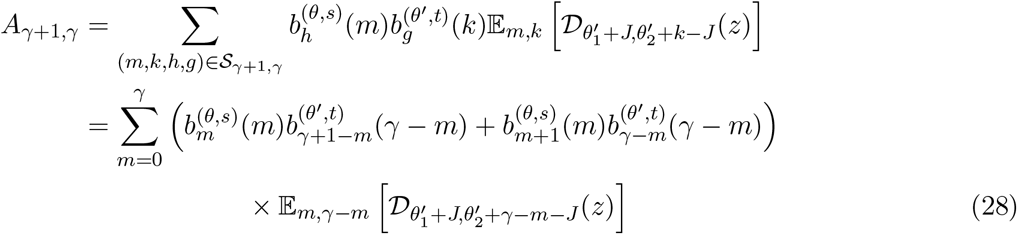

since *γ* + 1 − *h* = *g* ≥ *k* = *γ* − *m*.

In view of Lemma 2, we have that for 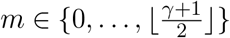

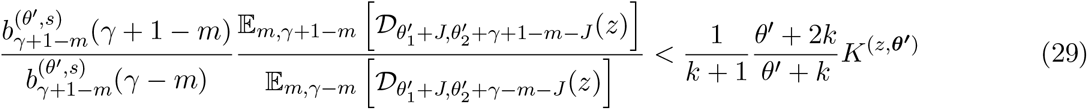

and that for 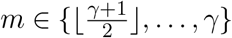

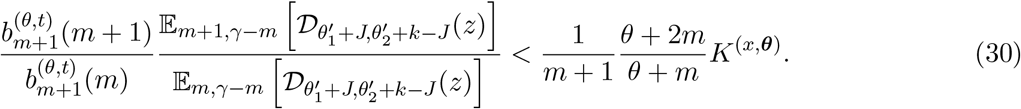

where we recall that *k* = *γ* − *m*. Observe that for 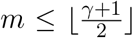, we also have that 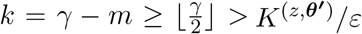, such that for the RHS of (29) we have

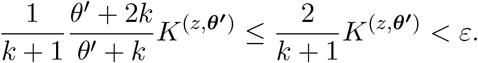

Similarly, for (30), 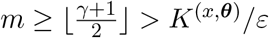, such that

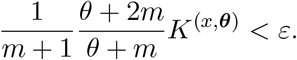

Thus (26) follows since for 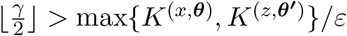, we have that

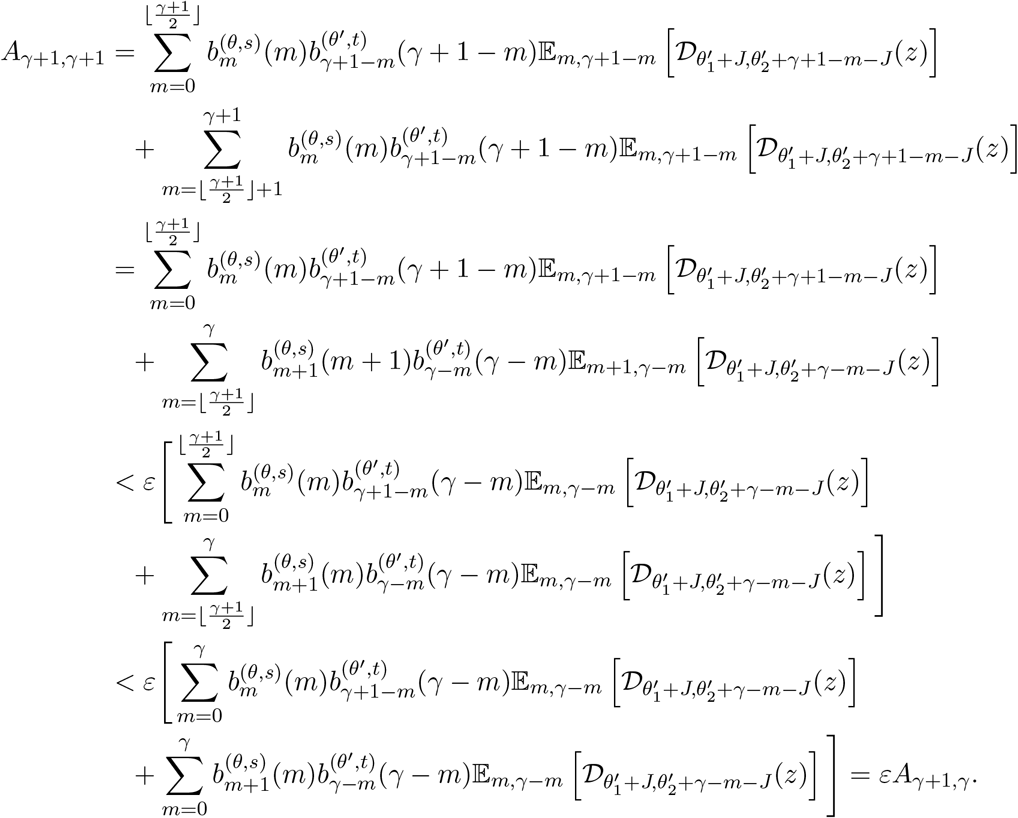

### Lemma 2.

*Let K*^(*x*,***θ***)^, *K*^(*z*,***θ ′***)^ *be defined as above. Then for any m* ∈ {0, …, *γ*} *and k* = *γ* − *m we have that*

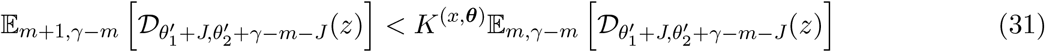

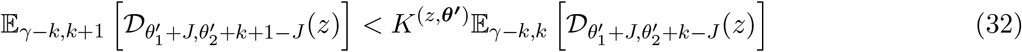

*Proof*. Assume that *l* ≤ ⌊*mx*⌋, and observe that

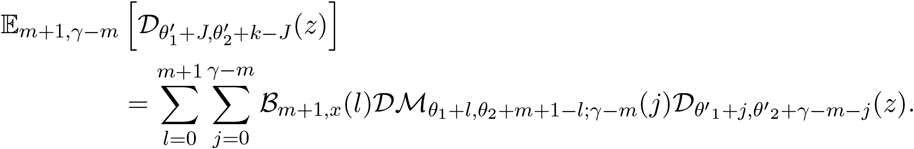

Additionally

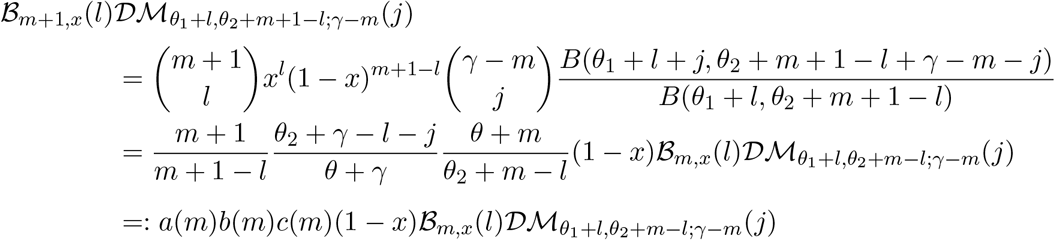

Trivially, *θ*_2_ + *γ* − *l* − *j* ≤ *θ*_2_ + *γ* ≤ *θ* + *γ*, so *b*(*m*) ≤ 1 independently of *m*.

For *a*(*m*), observe that this function is increasing in *m* such that for any *m* it holds that

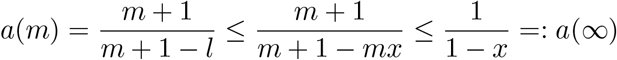

For *c*(*m*), we have that for *m* ≥ 1

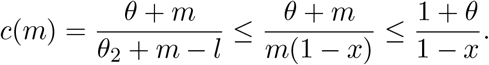

Putting all this together

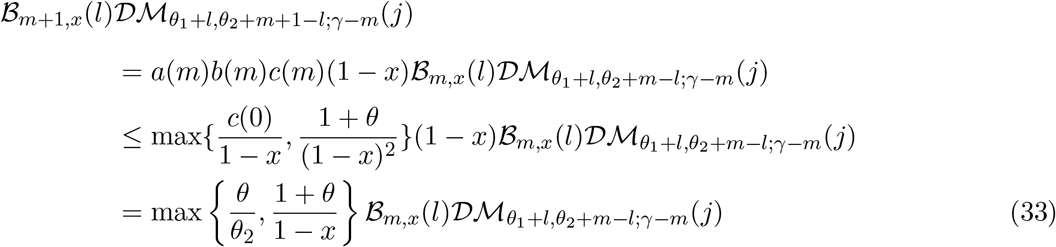

Summing from *j* = 0, …, *γ* − *m* and *l* = 0, …, ⌊*mx*⌋ we get

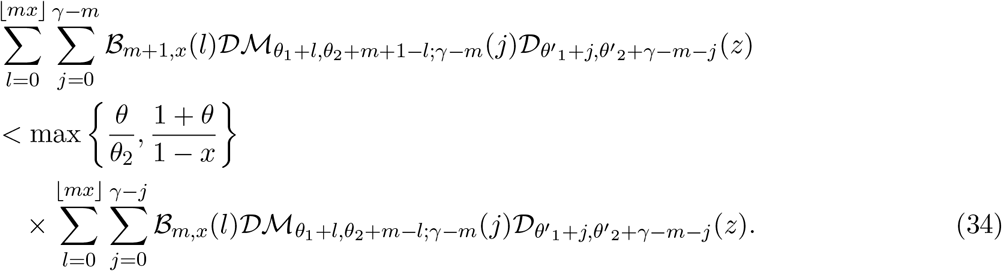

Suppose now that *l* ≥ ⌊*mx*⌋, then

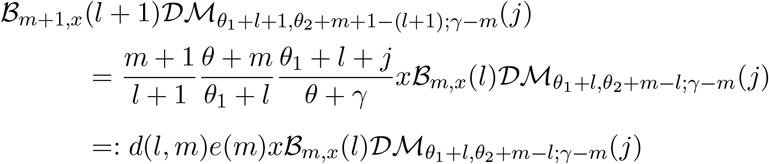

For *e*(*m*) we again have that trivially *θ*_1_ + *l* + *j* ≤ *θ*_1_ + *γ* ≤ *θ* + *γ*, so *e*(*m*) ≤ 1 independently of *m*, whilst for *d*(*l, m*), we have that if ⌊*mx*⌋ *>* 1, then

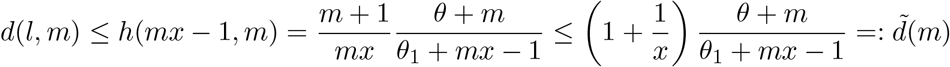

and (as in the display in the middle of p.27 in Jenkins and Spanò (2017)), since the sign of 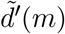 is independent of *m*, it follows that

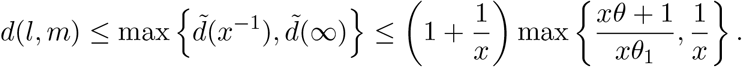

On the other hand, if ⌊*mx*⌋ ≤ 1, then

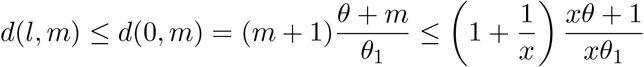

and thus it follows that independently of whether ⌊*mx*⌋ *>* 1 or ⌊*mx*⌋ ≤ 1,

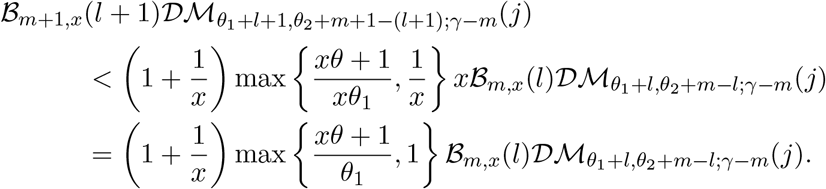

Summing over *j* = 0, …, *γ* − *m* and *l* = ⌊*mx*⌋ + 1, …, *m* + 1 we get that

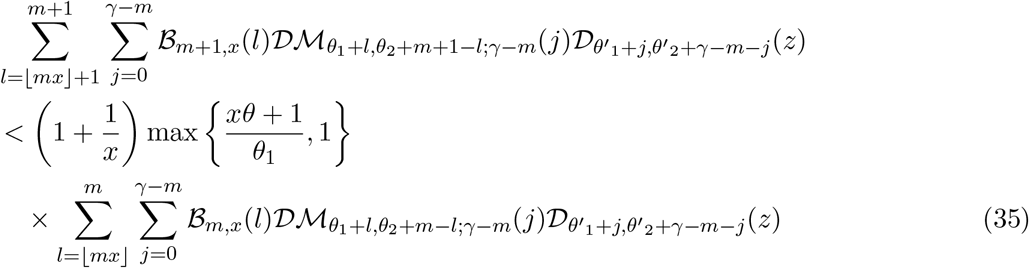

Summing (34) and (35) gives

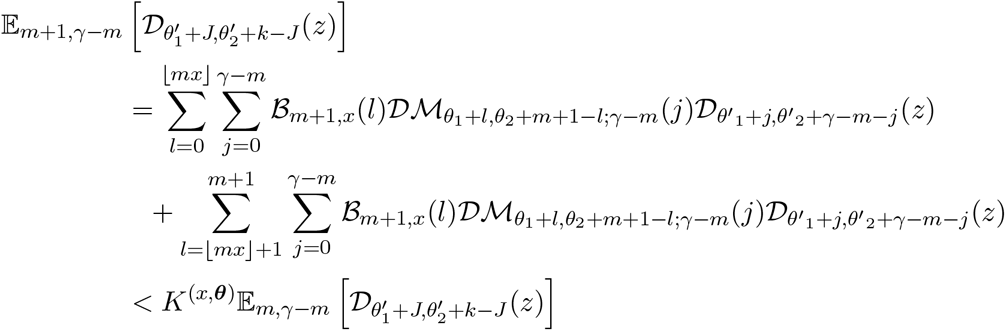

and thus (31) holds.

For (32), recall that *k* = *γ* − *m*, so that

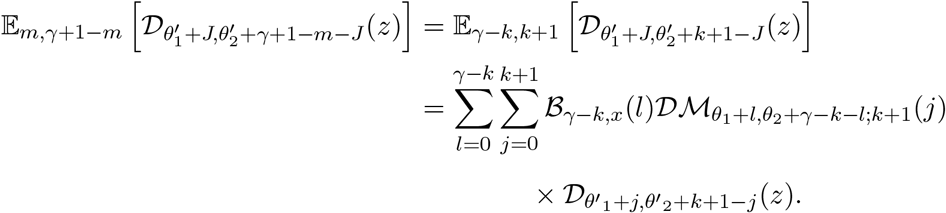

Suppose *j* ≤ ⌊*kz*⌋, then

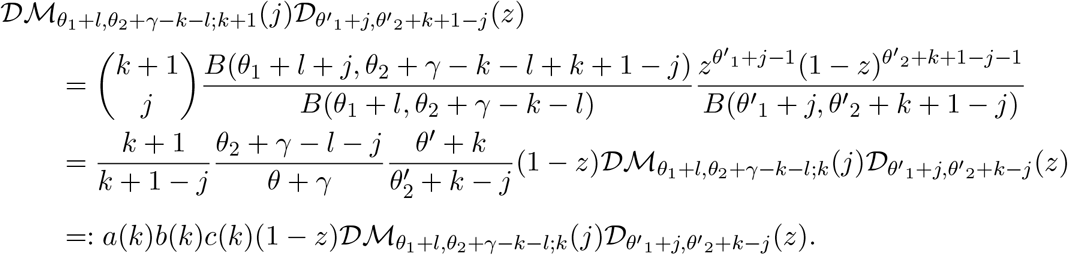

As done previously, we see that *a*(*k*) is increasing in *k* and furthermore

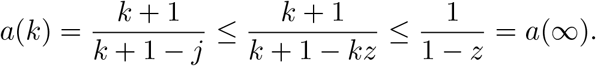

Also we have that *θ*_2_ + *γ* − *l* − *j* ≤ *θ*_2_ + *γ < θ* + *γ*, so that *b*(*k*) ≤ 1 independently of *k*.

For *c*(*k*) we once more have that for *k* ≥ 1

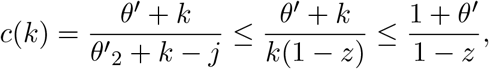

allowing us to deduce that

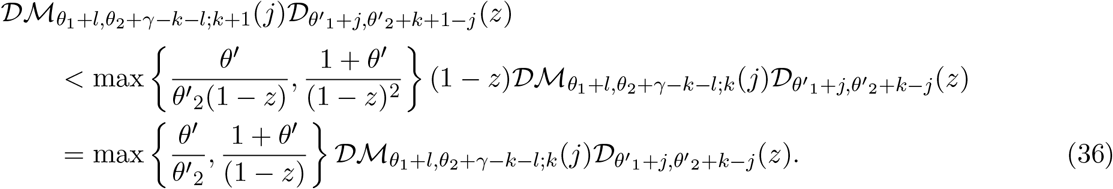

Thus, summing (36) over *j* = 0, …, ⌊*kz*⌋, and over *l* = 0, …, *γ* − *k* we get

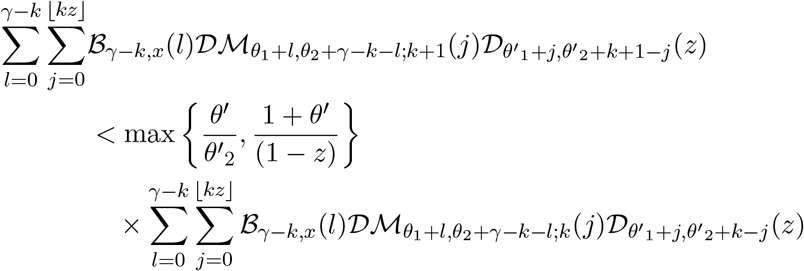

Now if *j* ≥ ⌊*kz*⌋, we have that

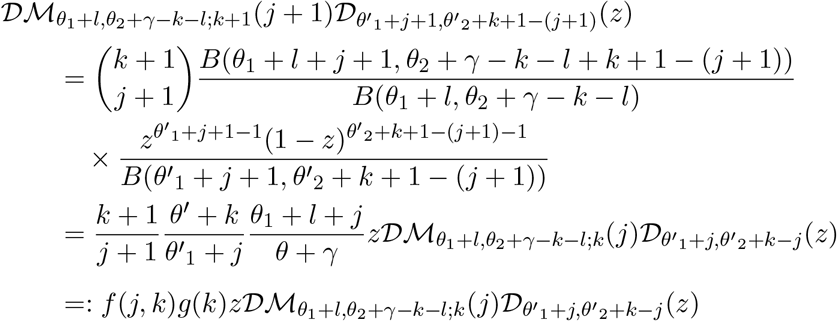

For *g*(*k*), recall that *j* ≤ *k* and *l* ≤ *γ* − *k*, so trivially *θ*_1_ + *l* + *j* ≤ *θ*_1_ + *γ* ≤ *θ* + *γ* and thus *g*(*k*) ≤ 1 independently of *k*.

For *f* (*j, k*), if ⌊*kz*⌋ *>* 1 we get that

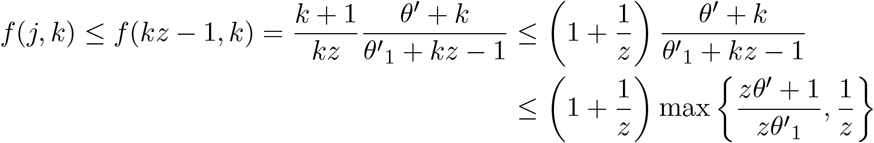

whilst if ⌊*kz*⌋ ≤ 1 it follows that

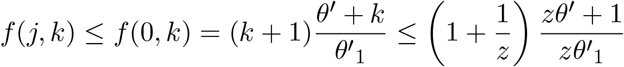

and thus regardless of the value of ⌊*kz*⌋ we have that

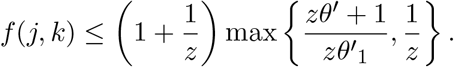

Thus

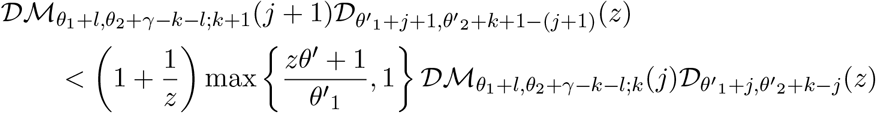

Summing over *j* = ⌊*kz*⌋ + 1, …, *k* + 1 and *l* = 0, …, *γ* – *k*

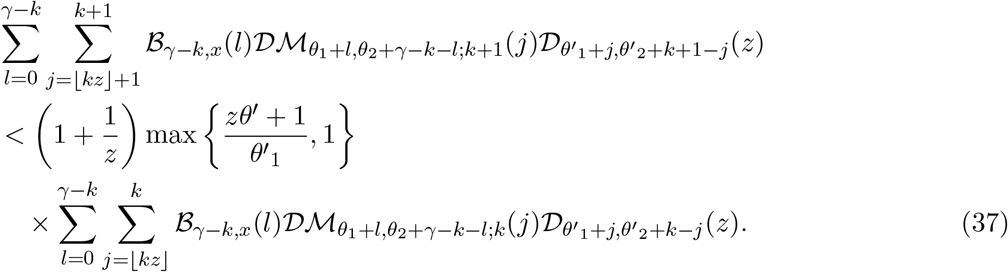

Summing (36) and (37) gives

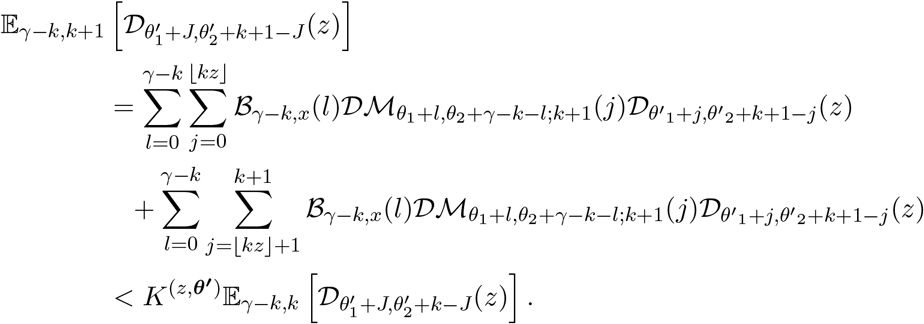

We can now use use Proposition 1 in Jenkins and Spanò (2017) and Proposition 1 above to deduce the following

### Proposition 3.

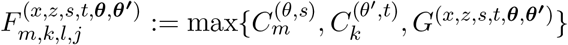 *and*

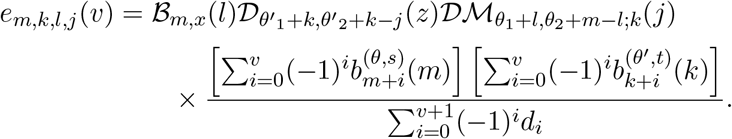

*Then for* 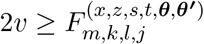 *it follows that*

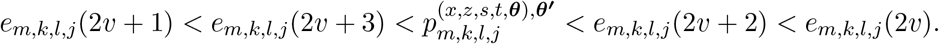

*Proof*. Follows immediately by applying Proposition 1 from Jenkins and Spanò (2017) to the numerator and Proposition 1 to the denominator.

## 3 Simulating draws from the transition densities (5)–(11) and (13)

For the remaining edge cases (i.e. where at least one of *x, z* is on the boundary {0, 1}), we can extend the same arguments made in Proposition 1 with some slight modifications to obtain eventually monotonic sequences of upper and lower bounds on the corresponding denominator. Sampling from the corresponding transition density then follows as usual. In particular, observe that if we can obtain such lower and upper bounds for the quantities

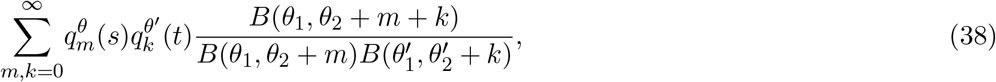

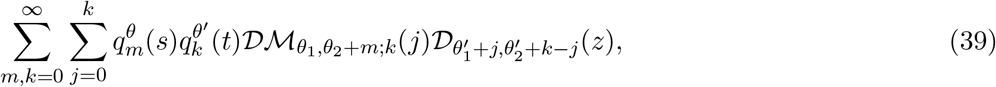

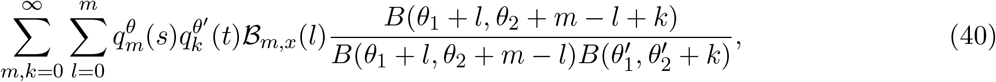

then the result follows for the remaining transition densities by considering suitably modified arguments.

We point out that in all three of the above cases, the exact same arguments in the proof of Proposition 1 can be used, with the only difference being that Lemma 2 needs to be suitably modified. To this end, it suffices to show that

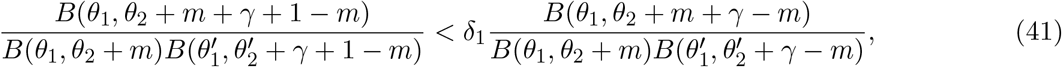

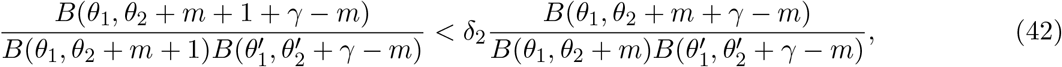

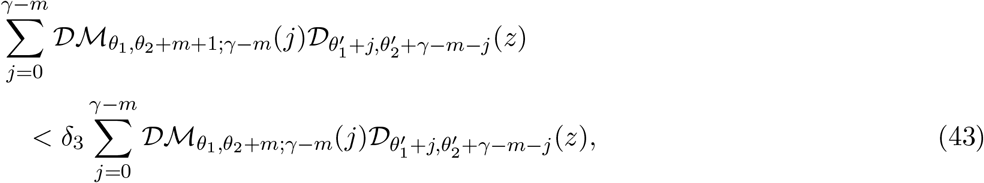

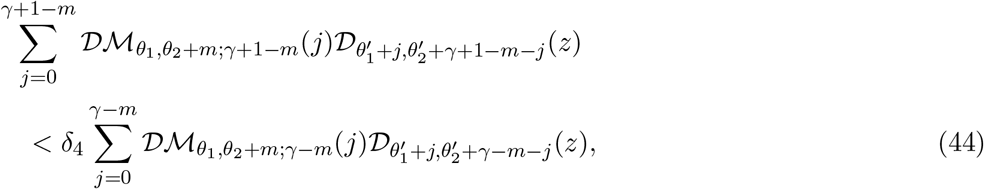

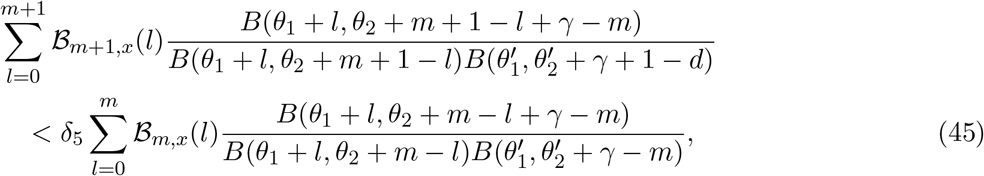

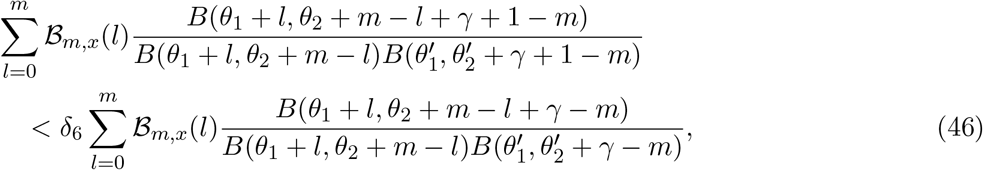

for an appropriate choice of *δ*_*i*_, with *i* ∈ {1, …, 6}. We start with (41) and (42) for which we observe that

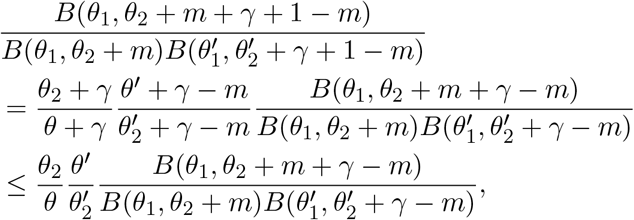

so 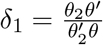, and

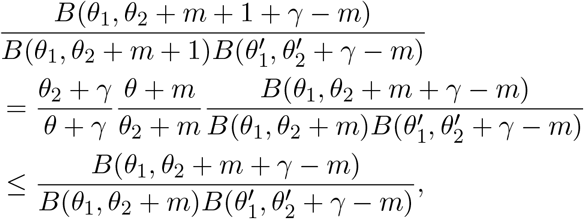

and thus *δ*_2_ = 1. Similar calculations give that

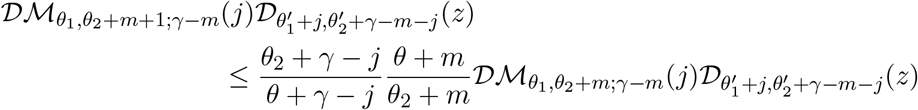

and thus summing over *j* = 0, …, *γ* − *m* gives that (43) holds with *δ*_3_ = 1. Further,

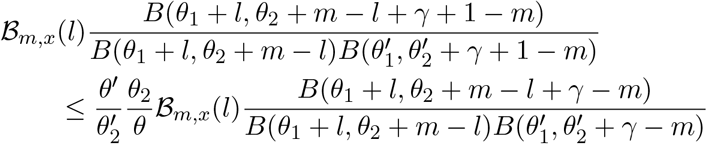

such that (46) holds with *δ*_6_ = *δ*_1_. The remaining bounds are slightly more involved but we can adopt the same approach as in Lemma 2, namely for (44) let *j* ≤ ⌊*kz*⌋, and then

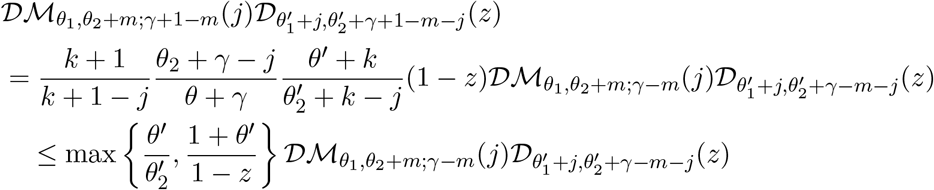

using the exact same arguments as in Lemma 2, and thus we can sum over *j* = 0, …, ⌊*kz*⌋ to get

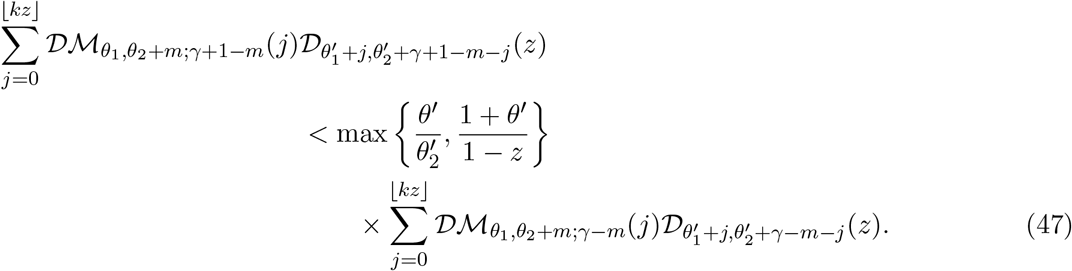

For *j* ≥ ⌊*kz*⌋ we get that

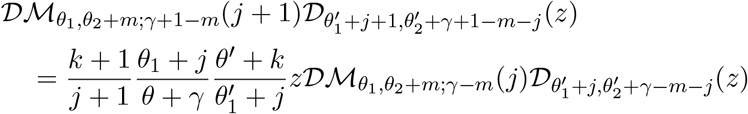

and again the same bounds as in Lemma 2 apply, such that summing over *j* = ⌊*kz*⌋ + 1, …, *γ* + 1 − *m* gives

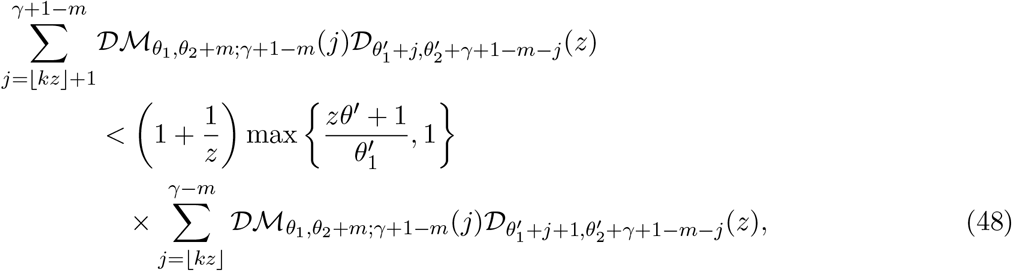

and summing (47) and (48) gives (44) with *δ*_4_ = *K*^(*z*,***θ′***)^.

We can repeat similar computations for (45), splitting over the case *l* ≤ ⌊*mx*⌋ and *l* ≥ ⌊*mx*⌋ to get the desired inequality with *δ*_5_ = *K*^(*x*,***θ***)^.

## 4 Small time approximations

For small times *s* or *t* − *s*, the quantities 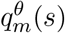 and 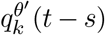 in (2) can become numerically unstable. To overcome this we make use of Gaussian approximations first developed in Griffiths (1984). We consider two specific cases:

1. When exactly one of *s* and *t* − *s* is less than some threshold *δ*
2. When both *s* and *t* − *s* are smaller than some threshold *δ*

In case 2, we employ Gaussian approximations for both 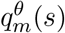 and 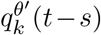 much in the same way as done in Sant et al. (2023). Case 1 however requires a separate approach as we observe that the decomposition introduced earlier making use of *A*_*γ,λ*_ relies on the joint decomposition of 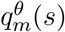 and 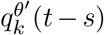, and thus if we approximate one by a Gaussian whilst we expand the other, we cannot recover the same scheme as in Proposition 1. Instead, assume for the moment that *s < δ < t* − *s*, then we compute the denominator by using the same decomposition as used in Proposition 4 in Jenkins and Spanò (2017), but where we now approximate the summation over *m, h* in (14) as follows

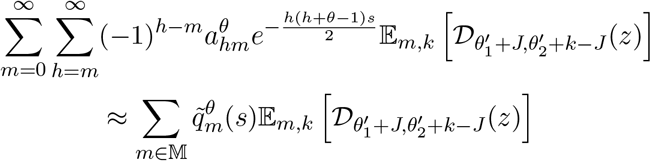

where 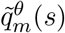 denotes the weight associated with the integer *m* under the discretised normal distribution used to approximate 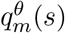, and M denotes the support of 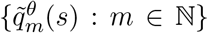 truncated to be within 5 standard deviations of the corresponding mean. The default setting in `EWF 2.0` is *δ* = 0.1 and can be tweaked by the user, however `EWF 2.0` features a robust safeguarding mechanism which automatically detects numerical instabilities (e.g. by detecting negative values for probabilities or lower bounds that exceed upper bounds) irrespective of choice of *δ*, falling back to Gaussian approximations whenever necessary.

## 5 Output verification

As in Section 7 of the Supplementary Information of Sant et al. (2023), we validate our implementation by comparing histograms of the draws obtained through the exact simulation algorithm to the empirical transition density evaluated by truncating the infinite series expressions for the corresponding bridge transition density. We consider two diffusion bridges, which we shall denote *B*1 and *B*2, with four epochs each. We summarise the mutation parameters, observation times and values, as well as sampling times we consider for both bridges in Table 2 and Figure 3, with the latter featuring an overlay of 100 candidate bridge paths for illustrative purposes.

**Table 2:**
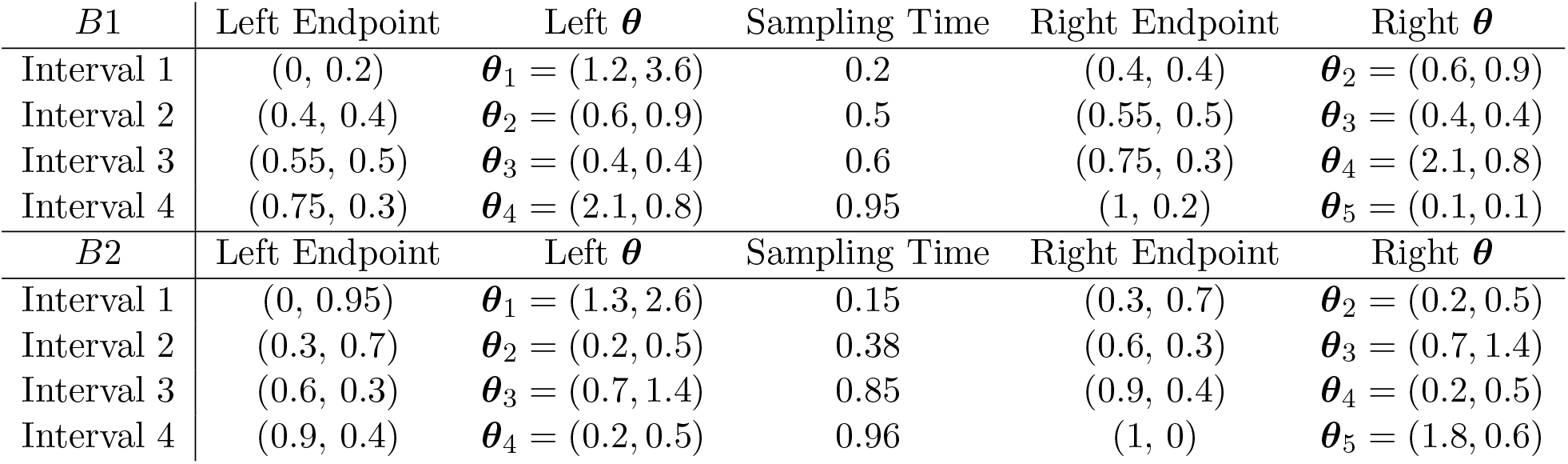
Sampling setup for bridges *B*1 and *B*2 where endpoints are reported as (*t, x*) where *t* is the time in diffusion units and *x* is the corresponding value of the diffusion bridge.

**Figure 3:**
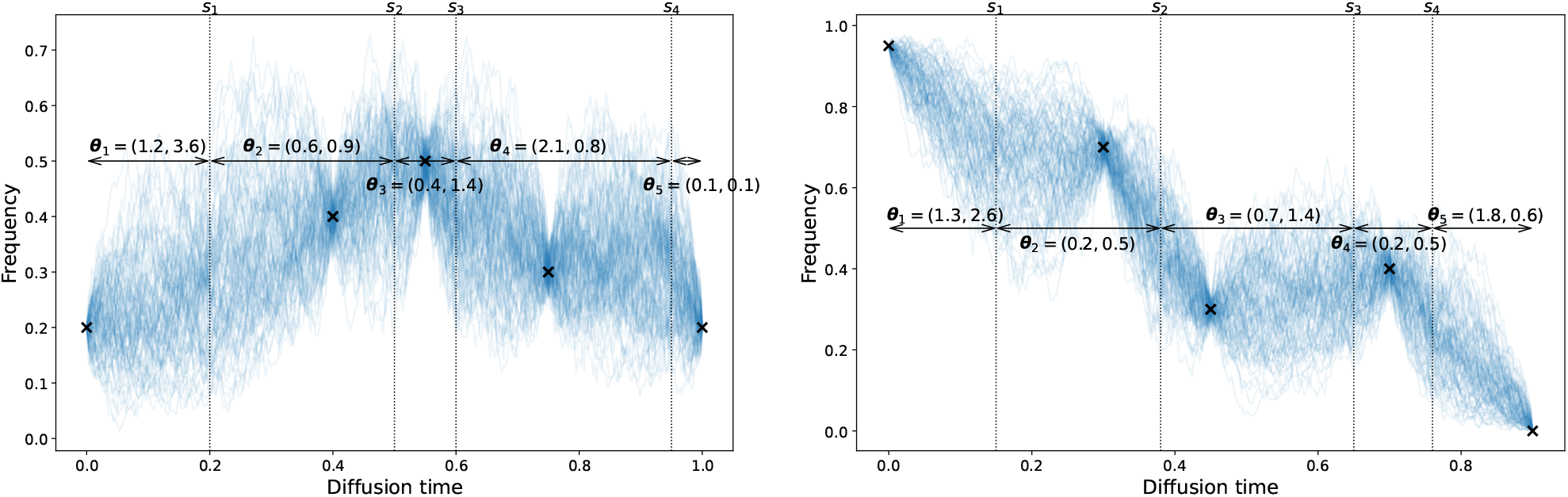
Sampling setups for bridges *B*1 (left) and *B*2 (right) with observations (black crosses), sampling times (vertical dotted lines) corresponding to times when the mutation parameter changes, and overlayed sample paths (blue). The mutation parameters adopted over each interval is annotated.

At each sampling time, we generate 10,000 draws from the corresponding law of the bridge process, and assess the performance of our implementation by comparing the resulting histogram to the empirical transition density, as well as QQ plots. Additionally we conduct a Kolmogorov–Smirnov goodness of fit test, where we compute the empirical cumulative density function through the empirical density function computed as a truncation of the corresponding infinite series expressions (2). As can be observed in Figures 4 and 5, our method targets the correct distributions in all cases considered.

**Figure 4:**
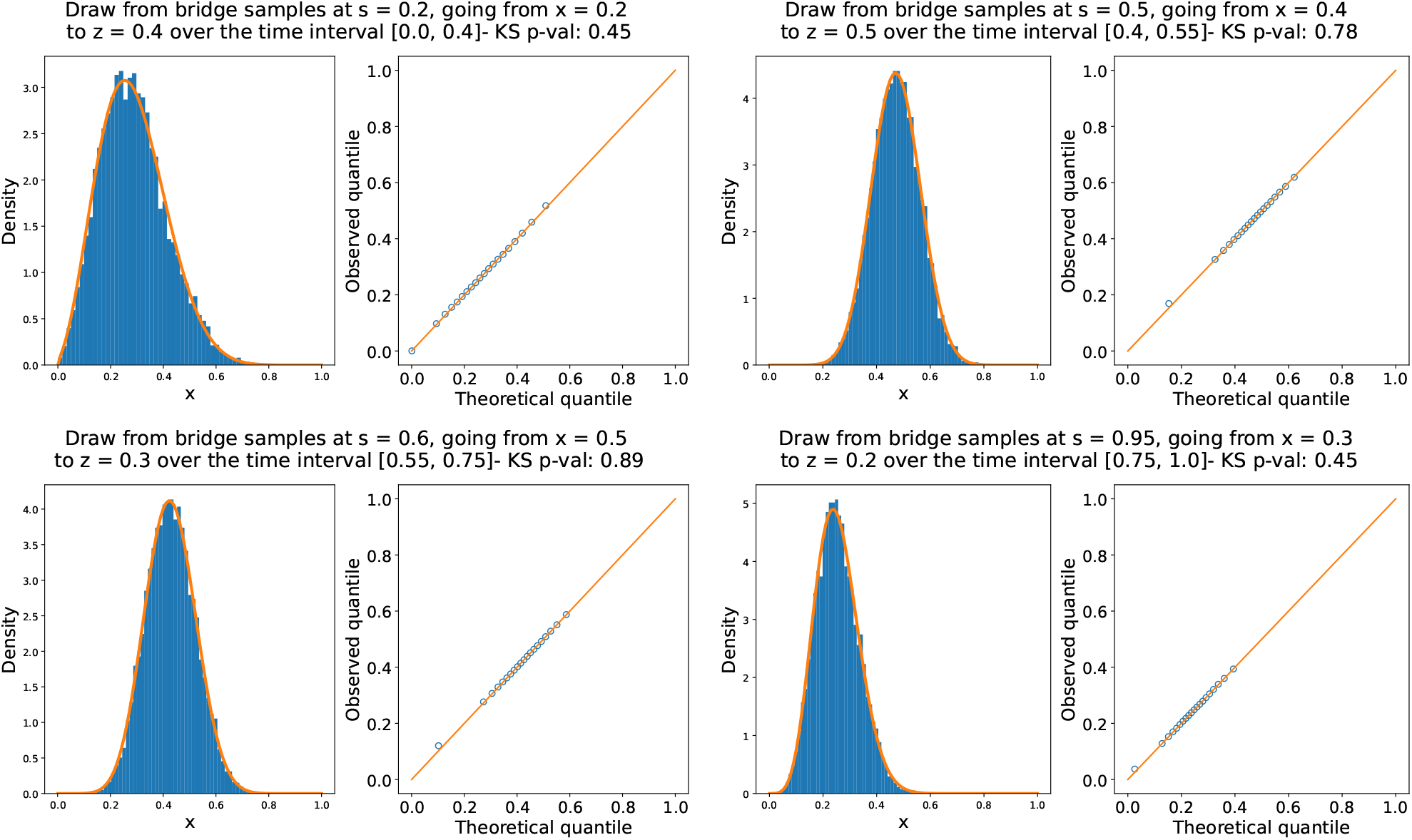
The output generated for bridge *B*1. Each panel corresponds to a different sampling time: *s* = 0.2 (top left), *s* = 0.5 (top right), *s* = 0.6 (bottom left), and *s* = 0.95 (bottom right). In each panel, we display the resulting histogram overlaid with the truncated density in orange on the left, whilst on the right we report the corresponding QQ plot. For visual clarity, the QQ plots display only 20 equally spaced empirical quantiles, although all goodness-of-fit statistics were computed using the full set of 10,000 simulated samples. The p-value obtained from the Kolmogorov–Smirnov test is reported in the title of each panel.

**Figure 5:**
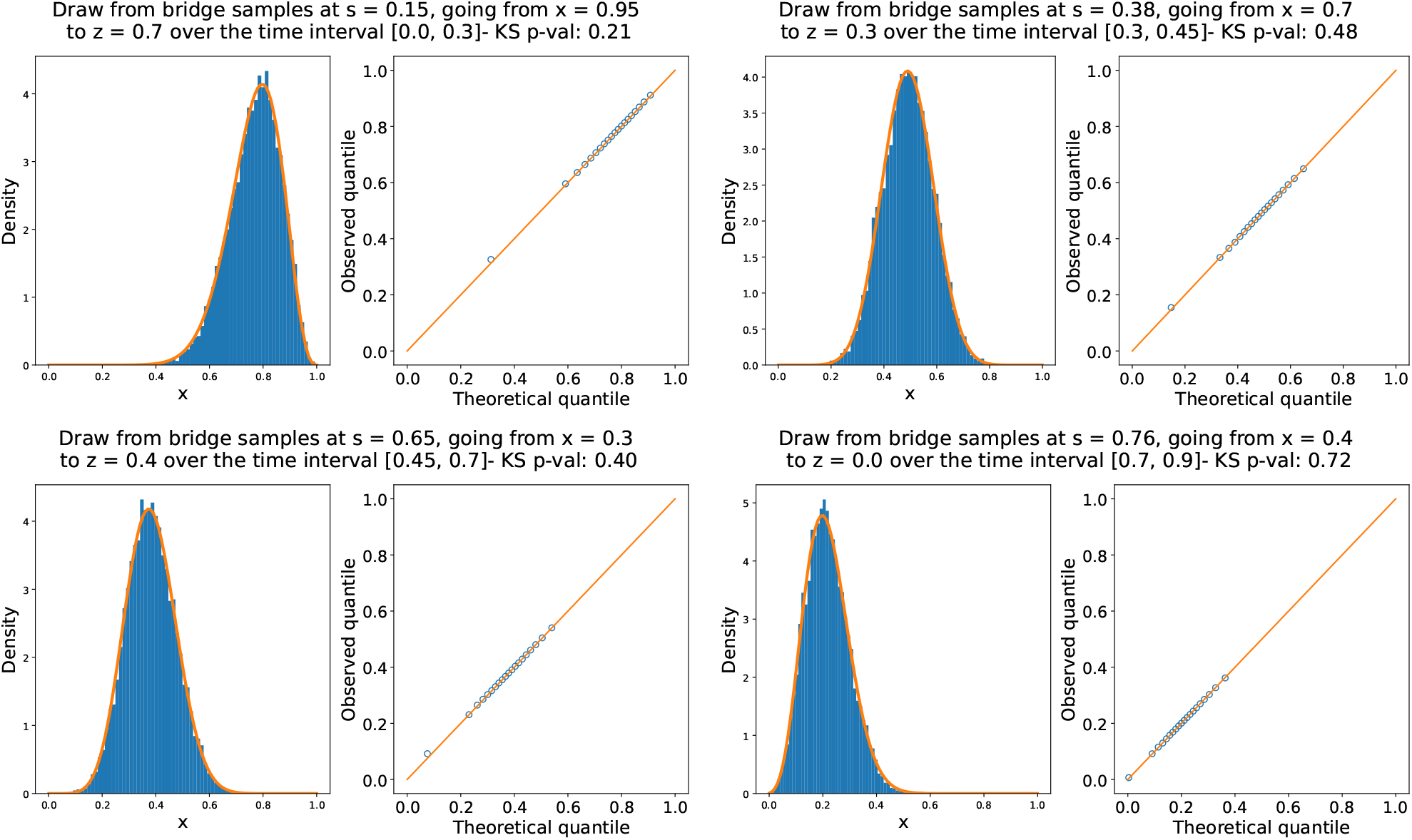
The output generated for bridge *B*2. Each panel corresponds to a different sampling time: *s* = 0.15 (top left), *s* = 0.38 (top right), *s* = 0.65 (bottom left), and *s* = 0.76 (bottom right). In each panel, we display the resulting histogram overlaid with the truncated density in orange on the left, whilst on the right we report the corresponding QQ plot. For visual clarity, the QQ plots display only 20 equally spaced empirical quantiles, although all goodness-of-fit statistics were computed using the full set of 10,000 simulated samples. The p-value obtained from the Kolmogorov–Smirnov test is reported in the title of each panel.

## 6 Non-neutral draws

In this section we give some more details on how a rejection sampler can be used to generate exact draws from the law of a non-neutral Wright–Fisher diffusion bridge. The crux is viewing the Radon–Nikodym derivative between the laws of neutral and non-neutral Wright–Fisher diffusions as (proportional to) the probability of an event that can be simulated; see (Jenkins and Spanò, 2017, Section 5) for further details. As in Jenkins and Spanò (2017), rejection efficiency decreases under increasingly strong selection, although the method remains exact across all parameter regimes.

Suppose now that apart from ***θ*** being the mutation parameter over [0, *s*), we also have that *σ* and ***η*** are the selection parameters, whilst over [*s, s* + *t*) we have ***θ***^***′***^, *σ*^′^ and ***η*** (observe that ***η*** remains the same over the interval [0, *s* + *t*)). Then denote by 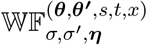 the corresponding path law of a Wright–Fisher diffusion started from *x*. By considering the marginal distribution of this diffusion at time *s* + *t* we get the decomposition

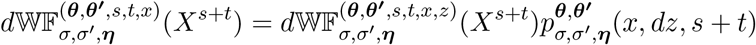

where 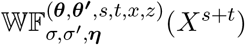 denotes the law of a Wright–Fisher diffusion *bridge* going from *x* at time 0 to *z* at time *s* + *t* and having mutation parameters ***θ*** over [0, *s*) and ***θ***^***′***^ over [*s, s* + *t*), as well as selection parameters *σ* over [0, *s*) and *σ*^′^ over [*s, s* + *t*). Furthermore, the Girsanov transform gives us that the Radon–Nikodym derivative between the laws of a neutral and non-neutral Wright–Fisher diffusion started from the same point *x* is

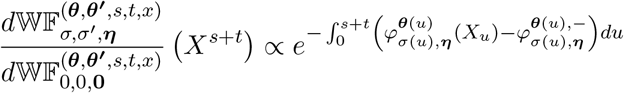

where

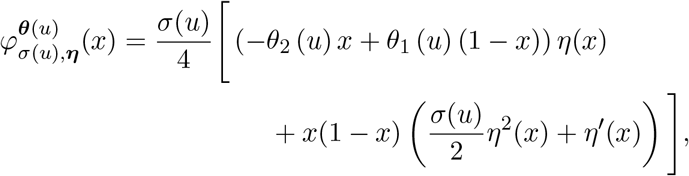

is a polynomial in *x* and thus can be bounded such that

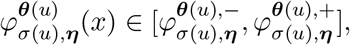

see (WF3) in (Jenkins and Spanò, 2017, Section 5.2). Thus we have that

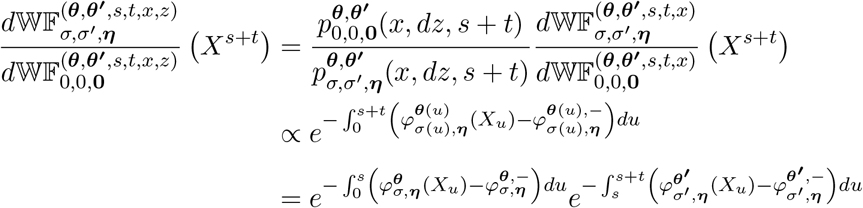

from which it follows that non-neutral bridges can be obtained by generating neutral candidate paths, and subsequently running a rejection step making use of two Poisson point processes, namely

- 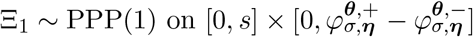
- 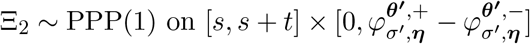

If the corresponding Poisson points all lie in the epigraph of 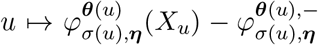, then the draw comes from the law of the non-neutral process as desired.

## 7 Demography for Figure 2

We use the demography inferred in Der Sarkissian et al. (2015) (as provided in Schraiber et al. (2016), specifically as found at https://github.com/Schraiber/selection/blob/master/horse.all.pop). In that file, time is given in diffusion units measured backward from the present, so *t* = 0 denotes the present and all observation times satisfy *t <* 0. Effective population sizes are scaled so that the most recent epoch has *N*_*e*_ = 1. Because our oldest reading corresponds to 20,000 years before present, we restrict our attention to the first two epochs of this demography; the second epoch extends well beyond our oldest observation and features an exponentially shrinking effective population size. Throughout, we index observation times as *t*_1_, *t*_2_, … with 0 *> t*_1_ *> t*_2_ *>* …, so in particular *t*_1_ *> t*_2_. `EWF 2.0` allows for piecewise constant demographies, and thus we need to truncate this demography and compute an appropriate effective population size with which to approximate the exponential curve over the shorter time interval we are interested in.

To this end, we write the effective population size as a function of time in diffusion units, namely

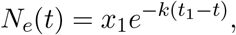

where *k* = (*t*_2_ − *t*_1_)^−1^(ln(*x*_2_) − ln(*x*_1_)) and we denote the effective population sizes in the first and second epoch as *x*_1_, *x*_2_ respectively, whilst *t*_1_, *t*_2_ denote the oldest time of the epoch (i.e. the time, in diffusion time units, at which there is a change in the population size). We shall henceforth reserve the notation *t* to denote time being measured in diffusion time scale, whilst *τ* is used to denote time as measured in years. Within the epoch having as left endpoint *t*_2_ and right endpoint *t*_1_, standard population genetics theory gives us that *dτ* = 2*gN*_*e*_(*t*)*dt*, where *g* is the generation time. Integrating and re-arranging using the above expression for the effective population size to get that for *t* ∈ [*t*_2_, *t*_1_] and *τ* ∈ [*τ*_2_, *τ*_1_]

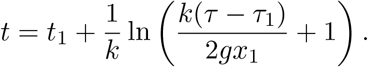

For the epoch we are interested in we have *t*_1_ = −0.0390625, *t*_2_ = −0.2109375, *τ*_1_ = 2*gN*_0_*t*_1_, *x*_1_ = *N*_0_, *x*_2_ = 6.875*N*_0_, and *τ* = −20, 000, which allows us to compute both *t* and *N*_*e*_(*t*) corresponding to the oldest observation recorded. Taking as initial population size *N*_0_ = 16, 000 and and generation gap *g* = 5 gives us the following demography, graphically summarised in Figure 6, and which was used to generate Figure 2 of the main text.

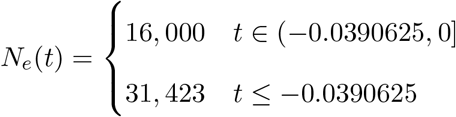

**Figure 6:**
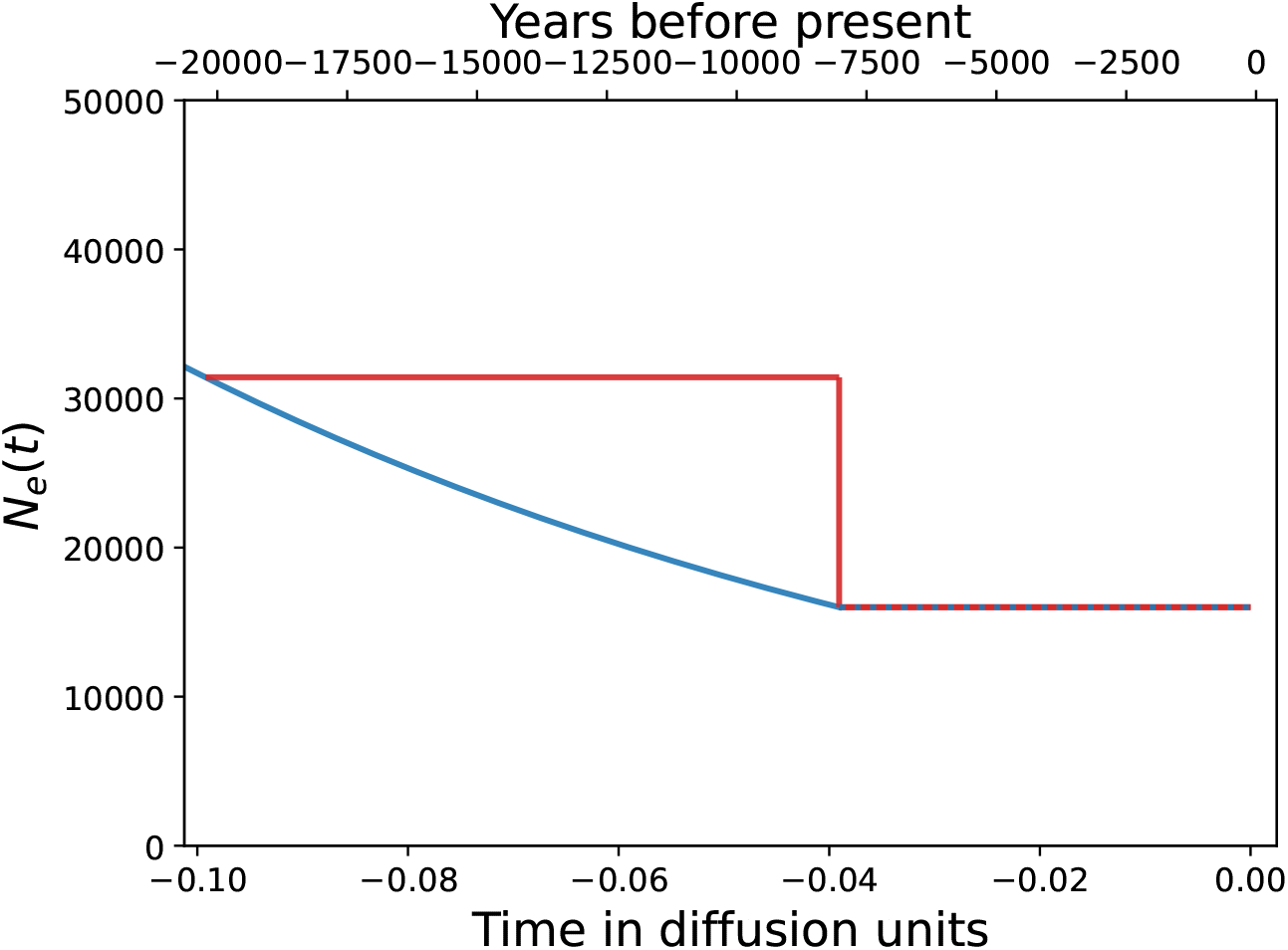
Plot of the demography inferred in Der Sarkissian et al. (2015) as provided in Schraiber *et al*. (2016) (blue), together with the piecewise linear approximation we make use of in Figure 2 (red).

## 8 Comparison with Wright–Fisher model simulations

To illustrate the computational advantages of the diffusion formulation, we compared the runtime of `EWF 2.0` with that of a standard discrete Wright–Fisher simulator. In the comparison, 10,000 neutral trajectories were generated under a constant population size, using mutation rates equal to those employed in the horse simulations in Figure 2. For each value of the effective population size *N*, the time horizon was held fixed in diffusion time units, ensuring that both methods were compared over the same evolutionary timescale.

The results are shown in Figure 7. As expected, the runtime of the discrete Wright–Fisher model increases approximately linearly with the effective population size. This reflects the fact that a fixed interval of diffusion time corresponds to *O*(*N* ) generations in the underlying discrete model, each of which must be simulated explicitly. By contrast, the runtime of `EWF 2.0` remains essentially constant across the range of population sizes considered. Note that the vertical axis in Supplementary Figure 7 is shown on a logarithmic scale, highlighting the substantial computational savings afforded by the diffusion-based approach for large populations.

**Figure 7:**
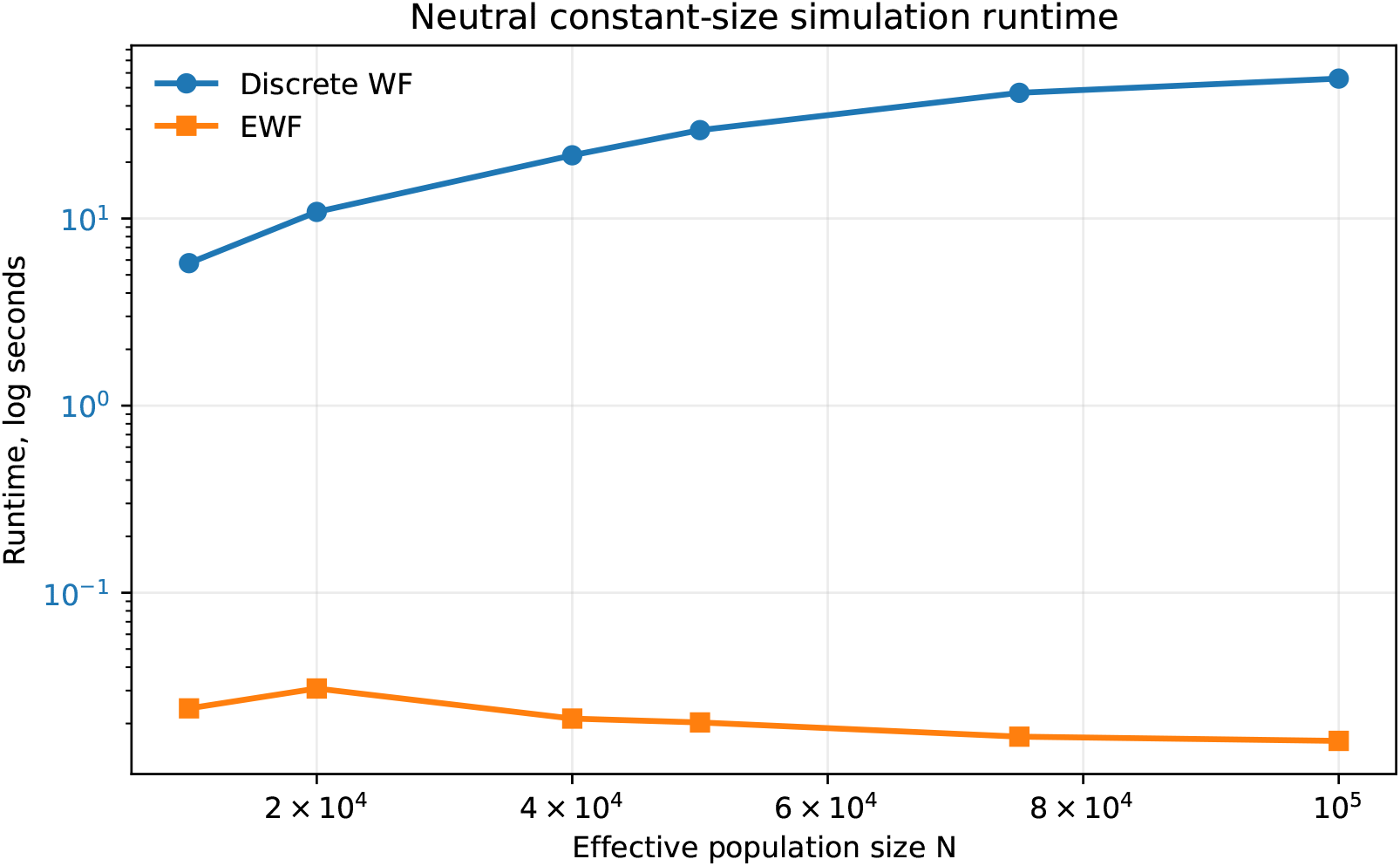
Run-time comparison to simulate 10,000 neutral samples when using a standard Wright–Fisher model (in blue) and `EWF 2.0` (in orange). Note that the *y*-axis is in log-scale, we fixed the time horizon to be 0.2 diffusion units, the mutation parameters to be *µ*_1_ = *µ*_2_ = 6.2 × 10^−9^, whilst varying the effective population size *N* ∈ {1, 2, 4, 5, 7.5, 10} × 10^4^.

